# FragLite mapping to identify the BRD4 recruitment site of P-TEFb

**DOI:** 10.64898/2026.04.09.717428

**Authors:** Ian Hope, Richard Heath, Arnaud Baslé, Mathew P. Martin, Michael J. Waring, Jane A. Endicott, Martin E.M. Noble, Natalie J. Tatum

## Abstract

The eukaryotic positive transcription elongation factor b (P-TEFb), composed of CDK9 and cyclin T, plays a central role in regulating RNA polymerase II (RNAPII). Phosphorylation of the RNAPII C-terminal domain (CTD) by P-TEFb promotes promoter proximal pause release and enables productive transcriptional elongation across many genes. Cyclin T mediates protein-protein interactions, several of which have been structurally characterised, that help to recruit and fine-tune P-TEFb activity to ensure a tight regulation of transcription. We have previously reported a set of halogenated chemical fragments termed FragLites that can prospectively identify protein interaction sites. Here, we report the FragLite map of cyclin T2, revealing binding sites corresponding to structurally defined cyclin T partners CDK9, AFF4, and HIV-1 Tat. Furthermore, we demonstrate the utility of FragLites in identifying a previously uncharacterised BRD4 binding site. By integrating FragLite clustering with biophysical analyses and AlphaFold3 modelling, we delineate the cyclin T-BRD4 interface. These analyses provide a comprehensive, chemically enriched fragment map highlighting functionally relevant sites to support future probe and modulator development to selectively target P-TEFb.

## Introduction

To maintain the protein complement of a cell, the activity of DNA-dependent RNA polymerase II (RNAPII) is tightly regulated [1] [2] [3] [4] [5]. The RNAPII C-terminal domain (CTD) consists (in human cells) of 52 evolutionary conserved heptad repeats and serves as a major site of transcription regulation [6] [7] [4] [8]. Subsequent to RNAPII recruitment to core promoters, the CTD undergoes multiple post-translational modifications to tune a stage-specific platform for the recruitment of mRNA processing enzymes, transcription regulatory factors and chromatin re-modelling proteins. Post transcription initiation (*circa* 20-50 nucleotides), RNAPII is recruited into an inhibitory complex containing DSIF (5,6-dichlorobenzimidazole 1-β-D-ribofuranoside [DRB]-sensitivity-inducing factor) [9] and NELF (negative elongation factor) [10] and undergoes a characteristic promoter proximal pause [11] [12] [13]. As a result, RNAPII pausing has been proposed to function as a regulatory checkpoint that promotes the efficient capping (and hence stabilisation) of RNA [14] [15] [16].

Failure to relieve pausing frequently leads to premature transcription termination [17] [18]. The positive transcription elongation factor b (P-TEFb), composed of CDK9 and cyclin T subunits, is the principal regulator of pause release [19] [20] [21]. P-TEFb phosphorylates DSIF and NELF, thereby promoting dissociation of inhibitory factors and enabling elongation [10] [22] [23] [24]. In addition, P-TEFb cooperates with CDK12-cyclin K to catalyse phosphorylation of Ser2 and Ser5 residues within the RNAPII CTD, further facilitating productive transcription [2] [25] [26]. Subsequently P-TEFb is incorporated into multiple alternative elongation complexes, for example through binding AFF1/4 into Super Elongation Complexes (SECs) that locate P-TEFb to promoters and enhancers [27].

In cells, the majority of P-TEFb is sequestered in an inactive state within the 7SK small nuclear ribonucleoprotein complex, which comprises 7SK snRNA [28] [29] [30] [31], HEXIM1/2, LARP7 and MePCE [32] [33] [34] [35]. Regulated release of P-TEFb from this complex and its delivery to promoter-proximal regions are essential for transcriptional activation. BRD4, a multi-domain protein of 1362 amino acids and a member of the BET bromodomain family, is capable of extracting and recruiting P-TEFb to sites of active transcription [36] [37] [38], but also has proposed P-TEFb recruitment-independent roles [39] [40]. It localises to active genes through its tandem bromodomains, which recognise acetylated lysine residues on histone tails, and through protein-protein interactions with transcription factors, Mediator complex components, and other regulators. The interaction between P-TEFb and BRD4 requires the C-terminal P-TEFb-interacting domain (PID) of BRD4 mapped to residues Thr1209-Phe1362 of which the extreme C-terminus (Arg1329-Phe1362) is required for P-TEFb binding [41] [42] [43] [44]. However, the precise molecular mechanisms controlling P-TEFb recruitment and release remain incompletely understood.

Beyond its canonical role in RNAPII-dependent transcription, BRD4 is hijacked by certain papillomaviruses to tether viral genomes to host chromatin during mitosis [45] [46]. This viral function also maps to the BRD4 C-terminal region, which can adopt a helical conformation upon binding partner proteins, as revealed in the structure of BRD4 bound to the human papillomavirus E2 protein [47]. These results suggest that the same structural element may represent a shared recognition motif in both viral and cellular interactions. Although CDK9 and cyclin T bind directly and independently to the PID, [41] the precise residues mediating these interactions have not been characterised.

The CDK9 and cyclin T subunits of P-TEFb include a protein kinase domain and a CBD containing tandem N- and C-terminal cyclin box folds (N-CBF and C-CBF) [48]. P-TEFb is distinguished by the splayed disposition of the CDK9 and cyclin T subunits that only interact through a relatively small interface (average buried surface area 974 Å^2^) involving their N-terminal lobes. This arrangement creates a prominent cleft between the C-terminal lobes that is either not present (for example in CDK2-cyclin A [49] or less prominent (for example in CDK4-cyclin D [50] [51]) or CDK12-cyclin K [52] in other CDK-cyclin pairs.

Cyclin T is distinguished from cell cycle-regulating cyclins (A, B, D and E) in that it lacks a canonical hydrophobic recruitment pocket within the N-CBF that recognises RXL motifs commonly found in CDK substrates and inhibitors ([53] [54] reviewed in [55]). Instead, structural studies show that protein recognition is mediated primarily through the C-CBF. The Human Immunodeficiency Virus (HIV-1) protein Tat redirects host transcriptional machinery to express genes required for virus replication [56] [57] [58] [59]. Tat engages the cyclin T1 C-CBF, but not cyclin T2 [60] [61] extending across the CBF interface to interact with CDK9 residues in the vicinity of the active site [62]. An N-terminal fragment of AFF4, a scaffolding component of the super-elongation complex folds upon binding to the cyclin T C-CBF including a site adjacent to that for Tat [63] [64] [65] [66].

We have previously described the use of FragLites for crystallographic screening, to identify and chemically map protein-protein interaction sites on CDK2, [67] CDK2-cyclin A [68] and together with PepLites to map interactions of the bromodomains of BRD4 and ATAD2 [69]. Protein-protein interactions are frequently mediated in part by docking of amino acid sidechains into poorly solvated, surface-accessible pockets of aromatic and non-polar groups which can make lipophilic and hydrogen bond interactions. FragLites are small compounds capable of emulating these native interactions. They offer two hydrogen bond forming substituents with differing spatial arrangements presented on an aromatic scaffold. When soaked at high (mM) concentrations into crystals of a protein target, binding events can be unambiguously detected aided by the inclusion of an anomalous scatterer within their structures. PepLites are molecules designed to recapitulate canonical amino acid sidechain contacts whilst retaining the intrinsic bromine anomalous scatterer.

Despite extensive functional studies, detailed structural information defining the BRD4 binding site on P-TEFb is lacking. Competition studies suggest that the binding of Tat and HEXIM1 to P-TEFb is mutually exclusive [70] [71] [72] [73]. BRD4 competes with HEXIM1 for cyclin T binding, [36] [74] implying that BRD4 may engage a region near the Tat site. BRD4 and AFF4 are proposed to form distinct active complexes with cyclin T, suggesting partially overlapping binding surfaces. However, the extended structure adopted by AFF4 on cyclin T does not reveal discrete interaction hotspots such as the recruitment pocket that might drive cyclin T association with this and other protein partners [63] [64] [65] [66]. We hypothesised that crystallographic FragLite screening of cyclin T would detect and probe hotspot sites that drive known cyclin T-AFF4 and -Tat interactions.

Here we present the FragLite map of cyclin T2. Well populated FragLite binding sites co-locate or are proximal to the binding sites of structurally characterised protein partners, and cluster on the C-CBF and at the N/C-CBF interface. We also detect one well populated site that does not co-locate with a known site of cyclin T-protein interaction and is in a region of high sequence conservation across both cyclin T orthologues. The FragLite map taken together with an AlphaFold multimer model of P-TEFb bound to BRD4 was used to generate a model of the cyclin T-BRD4 interface. This model informed the design of cyclin T mutants which we used to validate and characterise the cyclin T-BRD4 interaction *in vitro*.

## Results

Crystal structures of P-TEFb bound to its protein partners have been determined using cyclin T1. However, monomeric cyclin T2 is the orthologue that reproducibly crystallises to high resolution making it suitable for a fragment screen. Cyclin T1 and cyclin T2 exhibit a high degree of sequence conservation across the CBD (residues 1-281 that includes the N- and C-CBFs, 77% conserved, 84% similarity, (**Supplementary Figure 1**)), with divergence confined to their C-terminal regions.

We first determined the structure of CDK9-cyclin T2 (**Figure 1a and Figure 1d**). The structure confirms that the CDK9-cyclin T interface mediated by the N-terminal lobes of each protein and the mechanism of CDK9 activation are conserved between cyclin T orthologues (**Supplementary Figures 2a and 2b**). Phosphorylated CDK9 Thr186 within the activation segment (AS) is clearly resolved, forming direct and water-mediated contacts with conserved arginine residues (**Supplementary Figure 2c)**. Consistent with the mechanism of CDK activation as exemplified first by CDK2, cyclin T2 binding reorients the CDK9 αC (H1) helix to form a conserved Lys48-Glu66 salt bridge, rearranges the DFG motif (Asp167-Phe168-Gly169) at the start of the AS, and correctly aligns the catalytic triad (Asp149, Lys151, Asn154). Comparison with the structure of monomeric cyclin T2 (PDB 2IVX [48]) shows no major conformational changes in cyclin T2 upon CDK9 binding. The elongated Hc helix (Asn249-Ala262) is retained (**Supplementary Figure 2d**). From these structures we conclude that CDK9 activation by cyclin T is structurally conserved between orthologues and they support the use of cyclin T2 as a valid model system for crystallographic fragment screening to identify CDK9-cyclin T protein interaction sites.

**Figure 1.**
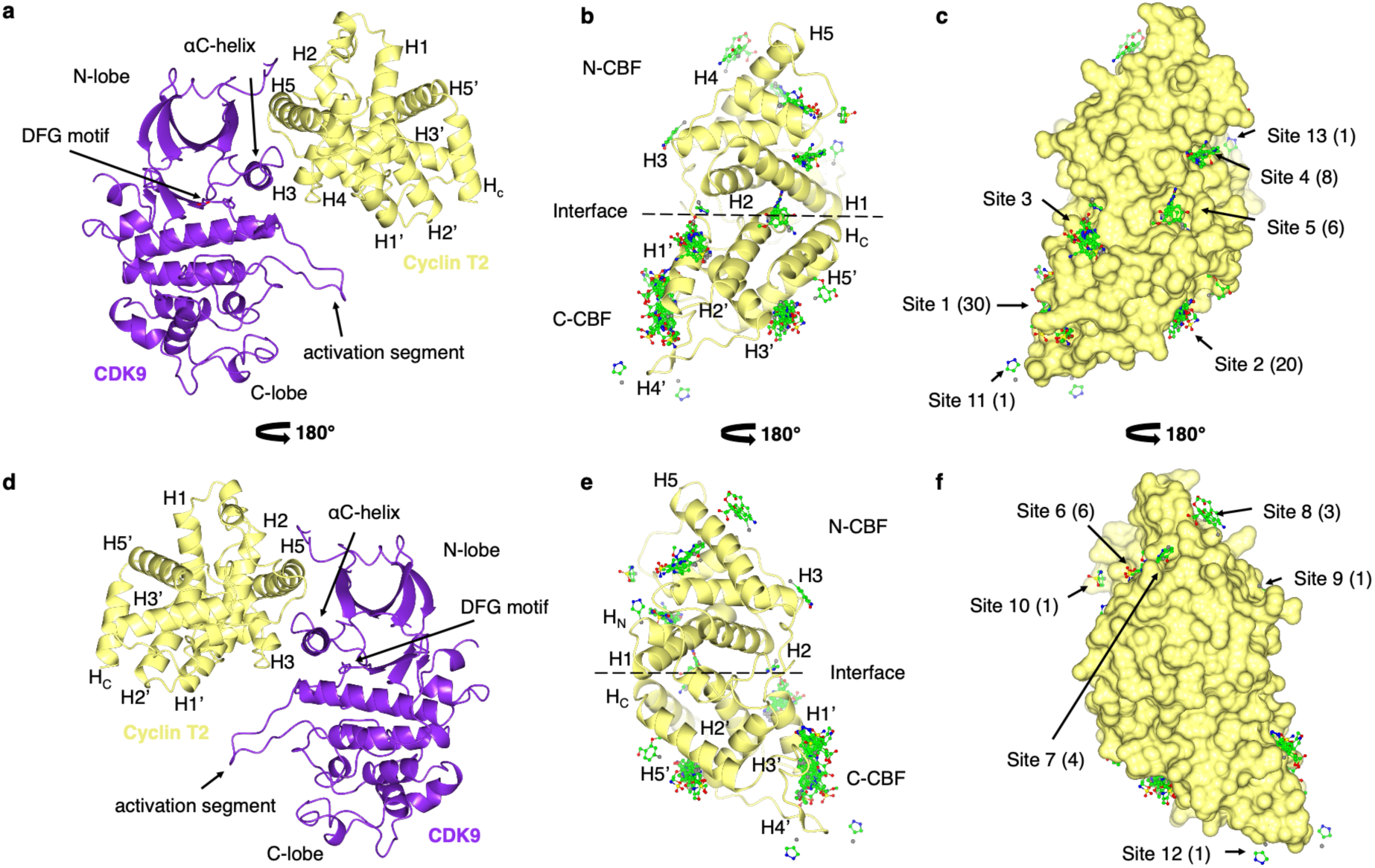
Structural characterisation of CDK9-cyclin T2 and cyclin T2. (a, **d)** Crystal structure of CDK9-cyclin T2. Key structural elements of CDK9 and cyclin T2 involved in cyclin-mediated activation of CDK9 are highlighted, including the CDK9 DFG motif (Asp167-Phe168-Gly169), αC-helix and activation segment. CDK9 and cyclin T2 are shown in magenta and lemon ribbon respectively. The cyclin T2 α-helices are labelled. (**b, e**) FragLite map of monomeric cyclin T2. The consolidated map generated by superposition of the individual cyclin T2-FragLite maps. There are significant clusters of FragLite binding with preferential enrichment of sites on the C-terminal Cyclin Box Fold (C-CBF). FragLite binders are coloured by heteroatom, with carbon atoms in green. (**c, f**) Molecular surface of cyclin T2. FragLites illustrate the fragment-accessible surface. The total number of binding events is 96, numbered by cluster. Seven sites (Sites 5-10 and 13) occupy the N-terminal Cyclin Box Fold (N-CBF), and six sites (Sites 1-4, 11 and 12) occupy the C-CBF. The sites are numbered based on the decreasing number of FragLite binding events detected (shown in brackets). (a) and (d), (b) and (e) and (c) and (f) are respectively related by a 180° rotation on the x-axis. Figure prepared using CCP4MG [95]. Related to Supplementary Figures 1, 2, 3, 4 and 5, Supplementary Tables 1, 2, 3 and 4 and validation reports supplied as supplemental material.

**Figure 2:**
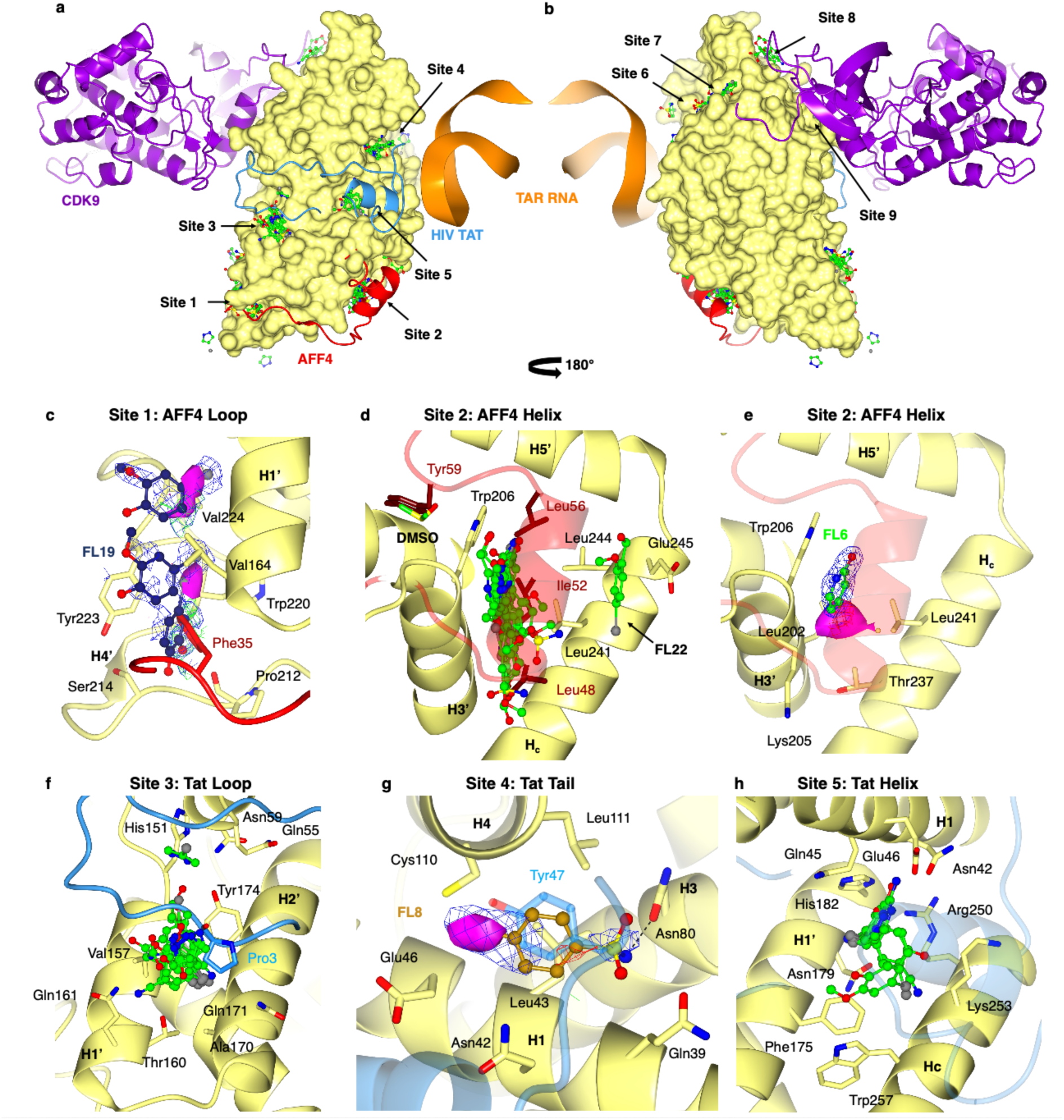
FragLite hotspots identify known protein-protein interaction sites. (**a, b**). Overlay of FragLite binding to monomeric cyclin T2 with P-TEFb protein partners derived from the crystal structure of CDK9-cyclinT1-AFF4-TAT-TAR (PDB 6CYT). CDK9 (purple ribbon), cyclin T2 (lemon surface), HIV-1 TAT (blue ribbon), AFF4 (red ribbon), TAR (orange ribbon). (**c**) Site 1, the “AFF4 Loop” as exemplified by FL19 (navy ball and stick). Anomalous difference density peaks (represented as solid magenta spheres) reveal multiple low-occupancy fragment orientations within the cleft predicted to accommodate AFF4 residues 22–33. Weighted 2Fo-Fc map is contoured at 0.3 e/Å³ (0.5 rmsd) and Fo-Fc difference map at ±0.6 e/Å³ (1 rmsd). FragLites at this site overlap the aromatic side chain of AFF4 Phe35 at the bottom of the cleft. (**d, e**) Site 2, the “AFF4 Helix”. Superposition of all fragment hits reveals a highly populated groove corresponding to the binding site of AFF4 residues 46–56. Fragments cluster over a surface formed by cyclin T2 residues Leu202, Lys205, Trp206 and Leu241. Structural overlay with the AFF4-bound complex (PDB 6CYT) shows spatial correspondence with the positions of AFF4 residues Leu48, Ile52 and Leu56. A DMSO molecule occupies the docking site of AFF4 residue Tyr59, occluding any additional interactions with Trp206. FragLites are shown in ball and stick representation, with carbon atoms in green and coloured by heteroatom. (**e**) Site 2 electron density map shown for FL6. The 2Fo-Fc weighted map contoured to 0.40 e/Å^3^, 1.48 rmsd and Fo-Fc difference map 0.31 e/Å^3^, 3 rmsd. (**f**) Site 3, the “Tat Loop” mapped on cyclin T2. All fragments (FL2, 4, 18, 19, 21, 22, and 23) converge with their bromine substituents into the shallow hydrophobic pocket occupied by the Tat Pro3 side chain near Tyr174. FL2 additionally occupies a second position within the site, with its hydrogen-bond donor-acceptor pair positioned adjacent to His151. (**g**) Site 4, the “Tat Tail” mapped on cyclin T2. FL8 (golden ball and stick) emulates the burial of Tat Tyr47 shown in blue. The FragLite aromatic core stacks against Leu43 and a hydrogen bond is formed between the NH_2_ of the sulfonamide group to Asn80 sidechain of cyclin T2. The weighted 2Fo-Fc map and Fo-Fc difference map both contoured to 0.34 e/Å^3^, 1.2 rmsd. Anomalous difference density peak for FragLite detection is represented as a solid magenta sphere. (**h**) FragLite binding at Site 5 the “Tat Helix” mapped on cyclin T2, showing fragments FL1, FL4, FL6, FL19, and FL21 overlapping a Tat helix (residues Cys27-Gly44, blue ribbon). All fragments converge on cyclin T2 Arg250, forming cation-π interactions with this cationic residue, which in cyclin T2 forms an electrostatic salt cluster with residues Asn42, Asn179 and His 182. Figure prepared using CCP4MG [95]. Related to Supplementary Figures 6, 7 and 8.

### FragLite mapping of cyclin T2

Members of the 31-member FragLite library [67] and 20-member PepLite library [69] (**Supplementary Table 1**), were soaked individually into single crystals of cyclin T2. Datasets having a maximal resolution between 1.6 and 2.4 Å (**Supplementary Table 2)** were initially processed using the XChem workflow [75] [76] and built-in Pan-Dataset Density Analysis (PanDDA) pipeline, [77] which identified 30 binding events from 12 individual FragLite crystal structures (**Supplementary Figure 3** and **Supplementary Table 3**). The more complex PepLite library provided six additional binding events across four ligand-bound crystal structures (**Supplementary Figure 4** and **Supplementary Table 3**). (PepLite nomenclature and structures are provided in **Supplementary Table 1**). Data were subsequently processed locally. Using an overlay of the anomalous log-likelihood gain (LLG) map with the 2Fo-Fc electron density map and Fo-Fc difference map, FragLite binding events were identified if the difference anomalous peaks were greater than four standard deviations above the mean value. Ninety-one binding events for 15 unique FragLite-bound structures were identified through providing the PanDDA map with the anomalous signal (**Figure 1b, 1c, 1e** and **1f**).

This analysis yielded a respectable hit rate of 48% (15/31) (**Supplementary Table 4**). Many of the ligands could be modelled with a high degree of confidence. However, in very low occupancy states, the presence of anomalous density taken together with non-spherical neighbouring electron density in accompanying electron density maps was deemed sufficient to support ligand binding (examples are given in **Supplementary Figure 5**). For FragLite 12, binding was assigned using the well-defined PanDDA event map electron density, despite a weak anomalous signal (3.88 rmsd) (**Supplementary Figure 5c**). The weak anomalous contribution is consistent with the reduced iodine signal at the data-collection wavelength. Including anomalous signal increased sensitivity, allowing us to detect 61 additional binding events and to resolve three more FragLite-bound structures than employing PanDDA alone.

Inspection of the FragLite map (**Figure 1b, 1c, 1e and 1f**) revealed there are 13 FragLite binding sites on the cyclin T2 CBD, seven and six respectively on the N- and C-CBFs. PepLites occupy three sites on the C-CBF overlapping the most heavily populated FragLite binding sites (Sites 1-3); and bind at a single site on the N-CBF not populated by the FragLites and only seen with PLThr and PLSer (Site 14, **Supplementary Figure 4**). Overall, the proportion of FragLite hits are predominantly clustered in sites on the C-CBF and at the CBF interface (74%, 67/90). Sites were selected for further analysis based on the number of FragLites bound and the diversity of interactions that the FragLites make. While individual fragments may sporadically bind to available surfaces and sites, an abundance of fragment binding is hypothesised to identify more meaningful functional hotspots. From this analysis we identified 6 sites (labelled Site 1- Site 6 in order of decreasing FragLite binding events) that had > 5 FragLites bound and > 6 binding events.

### FragLite binding identifies protein-protein interactions within P-TEFb complexes

We then analysed the distribution of binding events compared to the structure of cyclin T1 bound to AFF4, Tat, HIV-1 TAR (Trans-Activation Response RNA) and CDK9 (PDB 6CYT [78]) (**Figure 2**).

AFF4 is an intrinsically disordered protein that is proposed to fold upon binding to cyclin T. In all P-TEFb-AFF4 containing crystal structures (PDB 4IMY, [63] PDB 4OGR, [65] PDB 6CYT, [78] and PDB 4OR5 [64]) the AFF4 fragment comprising residues Asn2-Lys73 adopts an extended conformation in which 37 ordered residues (Pro33-Arg69) bind across the cyclin T1 C-CBF. This interaction defines the AFF4 helical region (residues Asp46-Ile66) and the loop region (residues Gln22-Glu45). Notably, residues Gln22-Ser32 remain unmodelled in all structures and are predicted to occupy the cleft between the splayed CDK9 and cyclin T subunits. One FragLite cluster (Site 1, the “AFF4 Loop” site, 30 binding events, **Figure 2c**) maps to the cleft region where the AFF4 loop is partially modelled (Pro33-Glu45). A second FragLite cluster is on the opposite face of cyclin T2 (Site 2, the “AFF4 Helix” site, 20 binding events, **Figure 2d and 2e**) and maps to a region that overlaps the first AFF4 helix binding site on cyclin T (Asp46-Ile56).

Site 1 is the most highly populated FragLite hotspot, highlighting its relevance for mapping the AFF4-CDK9-cyclin T interface. FragLites at this site overlap the aromatic side chain of AFF4 Phe35, which docks at the base of the site. However, the FragLites do not adopt a single defined pose. Instead, they produce a diffuse pattern of anomalous density (**Figure 2c**), indicative of multiple low-occupancy binding events rather than a stable binding mode.

We hypothesise that these signals, dispersed along the hydrophobic channel are structural substitutes for the flexible, unresolved portion of the AFF4 peptide (residues Gln22-Ser32), which are predicted to occupy the cleft between CDK9 and cyclin T.

The AFF4 helical region contains two tandem α-helices (residues Asp46-Gly57 and Tyr59-Ile66). Site 2 traces the groove accommodating the longer helix (residues Asp46-Ile56) (**Figure 2d and 2e**). Structural comparison with the AFF4-bound complex (PDB 6CYT [78]) shows that fragment binding recapitulates the hydrophobic interactions of AFF4 residues Leu48 and Ile52 with a cyclin T2 hydrophobic cluster contributed by Leu241 (cyclin T1 Leu242) and Leu244 (cyclin T1 Leu245), highlighting residue-level mimicry by the FragLites of the native interface. FragLite 22 (and PLGly, **Supplementary Figure 6a**) occupies the opposite face of this helix, revealing a complementary surface that mirrors the electrostatic character of the native interface, suggesting additional elements that may contribute to helical recognition.

Mutational studies underscore the functional importance of this site. Residues within or near the hotspot, most notably cyclin T1 Trp207 (cyclin T2 Trp206), are critical for cyclin T-AFF4 association [63]. One face of the site is capped by AFF4 Leu56, while the reverse face of Trp206 engages in a side-on π-stacking interaction with AFF4 Tyr59. The site partially overlaps a crystal lattice contact, and the reverse face of cyclin T2 Trp206 is further blocked by a DMSO molecule, a component from the fragment soak (**Figure 2d**). Although these factors may influence fragment positioning, they do not obscure the identification of the hotspot. Fragment mapping further identifies other residues, such as cyclin T2 Glu245 (cyclin T1 Glu246), that may contribute to recognition and represent previously underexplored features of this interaction surface.

FragLite binding also identifies protein interaction hotspots that drive HIV Tat binding. Tat, together with TAR RNA serves to recruit and activate CDK9-cyclin T1 (but not CDK9-cyclin T2), to enhance the transcription of HIV viral genes [56]. Tat, like AFF4, contains Intrinsically Disordered Regions (IDRs) that adopt an extended structure upon cyclin T1 binding, with the Tat-TAR recognition motif (TRM, cyclin T1 residues Arg254-Arg272, Hc helix) on the cyclin T1 C-CBF serving as the primary interaction site [79] [78] [65]. FragLite mapping of cyclin T2 identifies three distinct binding hotspots (Sites 3-5) that overlap structural elements known to mediate Tat interaction: the Tat N-terminal loop (Site 3, the “Tat Loop”, 13 binding events **(Figure 2f**)); the C-terminal tail (Site 4, the “Tat Tail”, eight binding events, (**Figure 2g**)) and the C-terminal helix (Site 5, the “Tat Helix”, six binding events, (**Figure 2h**)).

The N-terminal loop of Tat (residues 1-17) makes limited contacts with cyclin T1, driven largely by burial of hydrophobic side chains, and exemplified by a proline-aromatic C-H/π interaction between Tat Pro3 and a conserved tyrosine (cyclin T1 Tyr175/cyclin T2 Tyr174) (**Figure 2f**). Adjacent to the Tat Loop binding site, FragLites show rich occupancy (Site 3, the “Tat Loop” site), exhibiting a uniform burial of their bromine substituents into this pocket. FragLites exploit extensive opportunities within this cleft for hydrophobic and polar interactions as exemplified by FL22 which engages Tyr174 through a displaced π-π stacking interaction and extended water-mediated hydrogen bonds as well as hydrophobic interactions with Val157 (**Supplementary Figure 6b**). PepLite PLTyr also engages this site (**Supplementary Figure 6c**). This analysis particularly highlights the conserved tyrosine suggesting that the Tat protein might be repurposing in part an authentic protein interaction site. Mutagenesis studies demonstrate that substitution of cyclin T1 Tyr175 (Tyr175Glu, Tyr175Ser) severely impairs Tat binding, confirming the functional importance of this residue [73].

The intrinsically disordered Tat tail engages a flexible surface on cyclin T1 (Site 4, the “Tat Tail”). FragLites at this site adopt one or two binding poses (8 binding events). One orientation mirrors the burial of Tat Tyr47 and, in the case of FL8, recapitulates the Tat peptide hydrogen-bonding interaction with cyclin T1 Asn81 (cyclin T2 Asn80) (**Figure 2f**). In the alternative pose (as demonstrated by FL4, FL8 and FL21, **Supplementary Figure 6d**) hydrogen-bonding substituents engage cyclin T2 Glu46 (cyclin T1 Asp47) and Arg50 (cyclin T1 Arg51), reflecting the interactions formed by the Tat Tyr47 hydroxyl group in the Tat-cyclin T1 complex.

The C-terminal Tat tail contacts the cyclin T2 structural equivalent of the MRAIL helix (single letter amino acid code, helix H1) found in the N-CBF of canonical cell cycle cyclins. This sequence contributes to the hydrophobic recruitment site which accommodates flexible, non-helical peptide motifs (the RXL motif), found in various cell cycle CDK-cyclin substrates and inhibitors such as p27KIP1 [80] [81] (**Supplementary Figure 7a to 7c**). A structural comparison of cyclin T1 and T2 with cyclin A (**Supplementary Figure 7d to 7i**) illustrates that, while the overall surface topology is conserved, the chemical properties differ. Cyclin T1 and cyclin A present predominantly hydrophobic patches, whereas cyclin T2 displays a positively charged surface. These observations suggest that although Site 4 may be conserved between cyclin T isoforms it identifies a docking patch capable of accommodating diverse regulatory motifs, including IDRs, in a manner analogous to canonical cyclin-peptide interactions.

Site 5 is occupied by FL1, 4, 6, 19 and 21 and overlaps a third Tat binding region (the “Tat helix”, **Figure 2g, Supplementary Figure 6e)**. In the Tat bound cyclin T1 complex, cyclin T1 helices H1, H2’, and Hc create a critical interaction hub required for the folding of the Tat central helix (residues Cys27-Gly44). This folding is facilitated by Tat’s intramolecular coordination of a Zn ion and is reliant on the flexibility of the cyclin T1 Hc helix. Sequence divergence among residues within the Hc helix (**Supplementary Figure 1**) contributes to the orthologue-specific binding of cyclin T1 to Tat, highlighting structural differences between the two orthologues. Cyclin T2 displays a more rigid Hc helix while cyclin T1 exhibits flexibility. Selectivity is also determined by a critical coordinating residue within the TRM, Cys261 in cyclin T1 versus Asn260 in cyclin T2 [82] [83]. Mutations of Asn260 in cyclin T2 to cysteine facilitate Tat transactivation and suggest the cyclin T2 scaffold may also be amenable to Tat binding [84]. Given the requirement for secondary structure flexibility to distinguish Tat binding to the cyclin T orthologues, we are unable to correlate overlapping interactions made by FragLites with the Tat helix. However, the FragLites occupying Site 5 converge on Arg250, a cationic residue that in cyclin T2 forms stabilising ionic interactions with surrounding residues. The FragLites interact with Arg250 through cation-π stacking, highlighting a chemically favourable surface that may serve as a binding site for an authentic host partner. In cyclin T1, this residue (Arg251) is part of the TRM and contributes to TAR RNA coordination. FragLite richness and the favourable binding interactions at this site suggests the location of a binding site for an authentic host partner that has been re-purposed by the viral protein.

Collectively, the three FragLite-occupied sites that map to Tat binding regions display features consistent with protein-protein interaction hotspots. Within the extended Tat-interacting sequence, these sites may serve as focal points for stabilising contacts, highlighting surfaces that could be engaged by endogenous host proteins and that have been repurposed by the viral Tat IDR.

The next most populated FragLite binding site (Site 6, 6 binding events **Supplementary Figure 6f**) is located on the N-CBF within a deep pocket formed by helices H1, H3 and H4. It does not correspond to any currently annotated protein-protein interaction site on cyclin T, suggesting it may represent either an accessible interaction surface, or an uncharacterised protein interaction site. Cross-comparison with both cell cycle and transcriptional CDK-cyclin complexes reveals no protein partners engaging this region [85].

The FragLite clusters that map to CDK9 binding sites on cyclin T provide a useful point of comparison. CDK9 contacts residues from cyclin T helices H3, H4 and H5 within the N-CBF. Site 7 (the “CDK9 N-terminus”, **Supplementary Figure 8a**), Site 8 (the “CDK9 β-sheets”, **Supplementary Figure 8b and 8c**) and Site 9 (the “CDK9 α-helix”, **Supplementary Figure 8d**) map to the CDK9 interface (**Supplementary Table 4**). Each site shows low fragment occupancy (4, 3 and 1 event respectively) and therefore do not emerge as FragLite hotspots under our selection criteria. Site 7, the most populous of these three FragLite binding sites binds FL2, FL4, FL21 and FL23 and is located at the interface between the H2-H3 linker and the H5 helix (**Supplementary Figure 8a**).

Taken together FragLite clusters define compact, energetically favourable pockets that are characteristic of PPIs mediated by multipartite, modular interaction sites. In this context, FragLites provide a complementary view of cyclin-mediated protein-protein interactions to that provided by the structures of cyclin-containing multiprotein assemblies. Notably, PepLites also overlap the most populated FragLite sites, providing independent support that these surfaces represent genuine, energetically favourable interaction hotspots. This framework enables the identification of fragment-enriched surfaces, including a highly populated pocket on the C-CBF (Site 3), which is examined further below.

### The cyclin T2 FragLite map identifies residues that drive the interaction between cyclin T and BRD4

Tat and BRD4 associate with P-TEFb in a mutually exclusive manner [41] suggesting that at least part of the BRD4 binding site may overlap with one of the FragLite clusters (Sites 1-3) that collocate with the Tat binding site. Among the Tat-overlapping hotspots, high FragLite occupancy marks Site 3 as a candidate endogenous protein-interaction hotspot. The site is fragment rich, not significantly influenced by crystal contacts and is not used as a PPI site by structurally characterised non-viral P-TEFb partners. Importantly, Site 3 lies closest to the CDK9-cyclin T2 interface, consistent with a binding mode in which a host regulatory protein might engage both subunits of P-TEFb [41]. While this hydrophobic patch partially overlaps the Tat interface in cyclin T1, its strong conservation across cyclin T orthologues suggests that it did not evolve for viral recognition. Instead, we hypothesised that Site 3 may correspond to a binding site for a host factor that competes with Tat.

In the absence of structural data for this interaction, we employed AF3 [86] to generate predicted models of P-TEFb, comprising both orthologues of cyclin T and the experimentally mapped PID interacting region of BRD4 (residues Thr1309-Phe1362) to explore plausible cyclin T-partner complexes at this hotspot (**Figure 3a and 3b, Supplementary Figure 9a and 9b**). This prediction reveals that BRD4 employs tandem amphipathic helices formed by residues Pro1317-Ala1342 (N-terminal helix) and Ala1349-Phe1362 (C-terminal helix) to bind to CDK9 and cyclin T2 respectively. The N-terminal helix of the pair is positioned adjacent to the C-lobe of CDK9 (**Figure 3a and 3b**). This tandem helical BRD4 motif was previously observed in the structure of BRD4 bound to human papillovirus E2 (PDB 2NNU [47]). The helices are connected through a linker region (residues Ala1342-Asn1348) that engages with the activation segment of CDK9.

**Figure 3.**
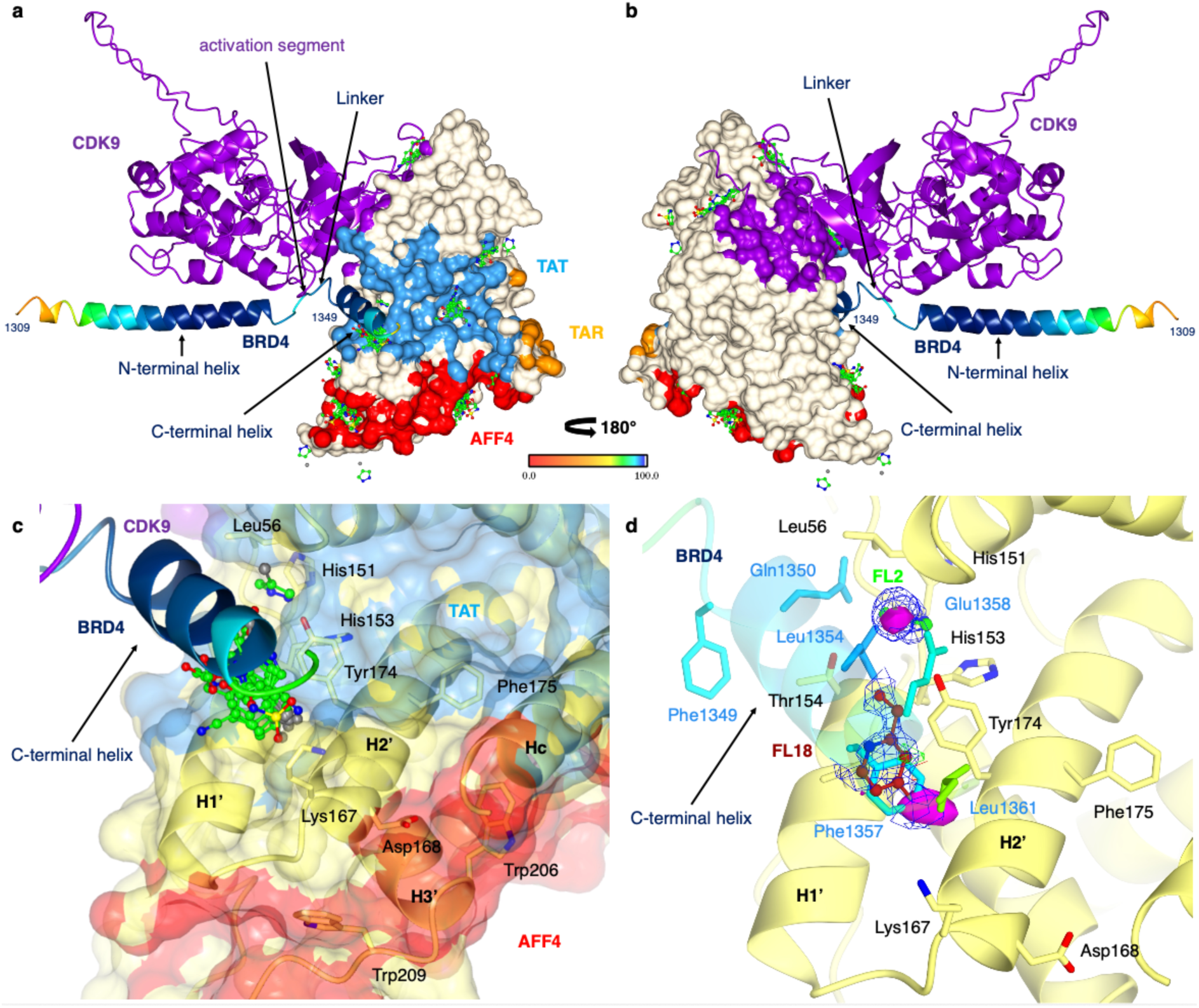
The AlphaFold multimer model of the BRD4 C-terminus with CDK9-cyclin T2. (**a, b**) BRD4 binding to cyclin CDK9-T1. BRD4 spans the C-lobe of CDK9 to contact the activation segment before intersecting the N/C-Cyclin Box Fold (CBF) interface of cyclin T1. The cyclin T1 surfaces bound by CDK9, AFF4, Tat and TAR in the structure of the pentameric complex that includes cyclin T1 (PDB 6CYT) are coloured purple, red, blue and orange respectively. The BRD4 model is coloured by the associated predicted Local Distance Difference Test (pLDDT) score [96]. Structures are rotated by 180°. (**c**) FragLites occupy a hydrophobic channel on cyclin T2 between the H1’ and H2’ helices of the C-CBF, overlapping the modelled binding site of the BRD4 C-terminal helix (residues Ala1349-Phe1362). (**d**) FragLite binding mimics interactions of three modelled BRD4 residues (Phe1357, Glu1358, and Leu1361), highlighting a potential hotspot for the BRD4 interaction. For example, FL2 (green) emulates Glu1358, while FL18 (maroon) aligns with the aromatic Phe/Leu side chains, with its aromatic core overlapping the phenylalanine sidechain and bromine substituent overlapping the leucine sidechain respectively. BRD4 model secondary structural elements and amino acid sidechains are coloured by the associated predicted Local Distance Difference Test (pLDDT) score [96]. The weighted 2Fo-Fc map and Fo-Fc difference map for FL2 and FL18 are contoured to 0.30 e/Å^3^, 1 rmsd and 0.34 e/Å^3^, 0.7 rmsd, respectively. FragLite anomalous difference peaks are shown as magenta spheres. Figure prepared using CCP4MG [95]. Related to Supplementary Figure 9 and Supplementary Figure 10.

A comparison of the AF3 model of the CDK9-cyclin T2-BRD4 complex with the cyclin T2 FragLite map reveals the convergence between the predicted BRD4 interface and FragLite Site 3 (the Tat Loop Site) (**Figure 3c**). FragLites emulate key interactions predicted for BRD4, including hydrophobic packing around cyclin T2 Leu56, His151, His153, and Val157, and aromatic or polar contacts involving Tyr174 and Thr154 (**Figure 3d**). This FragLite hotspot is distinct from the AFF4 binding site (**Figure 3a**) and the collocated FragLite sites (Site 1, the AFF4 Loop and Site 2 the AFF4 Helix) (**Figure 2a-2d**) indicating that it constitutes a distinct regulatory surface on cyclin T. The concordance between predicted interfaces and experimentally mapped fragment hotspots is consistent with Site 3 functioning as a BRD4-recognition surface and was the basis on which residues were selected for functional validation.

Notably, BRD4 has also been reported not to induce displacement of HEXIM1 from P-TEFb [37]. P-TEFb binds to HEXIM1 within an inhibitory particle that also includes 7SK snRNA, LARP7 and MePCE suggesting that displacement of P-TEFb from HEXIM1 might require more significant protein assembly re-organisation rather than direct HEXIM1-BRD4 competition for binding. To explore this possibility, we again used AF3 to generate a model of a ternary CDK9-cyclin T-HEXIM1 (residues Met255-Asp359) complex (**Supplementary Figure 10a and 10b**). This model positions one helix of HEXIM 1’s coiled coil motif between the cyclin T1

H1’-H2ʹ helices overlapping with both FragLite Site 3 (the Tat Loop Site) and the modelled BRD4 site (**Supplementary Figure 10c and 10d**). Mutagenesis experiments indicate that this site is also used by HEXIM1, as mutation of cyclin T1 Tyr175, in addition to Lys168 and Asp169 at the base of the H2’ helix, abolishes HEXIM1 binding [73] (**Supplementary Figure 10e and 10f).** The modelled HEXIM1 binding site on cyclin T is extensive and would also be incompatible with AFF4 binding [72] (**Supplementary Figure 10c and 10d**). The models suggest that the cyclin T HEXIM1 and BRD4 binding sites overlap in part and with a FragLite binding hotspot exploited by HIV Tat. It can be hypothesised that the H1’-H2’ channel forms a general docking site for regulatory motifs. Taken together, FragLite mapping, mutagenesis studies, and structural analysis support the existence of a common, chemically adaptable docking surface on cyclin T that accommodates both viral and host regulatory partners.

To experimentally confirm the proposed BRD4 binding mode, we first defined the minimal BRD4 fragment required for cyclin T interaction. The C-terminal sequence of BRD4 binds to CDK9 and extends across to cyclin T. Previous studies have shown that both CDK9 and cyclin T can independently interact with the C-terminal 34 amino acids of BRD4 (residues Arg1329-Phe1362) [41]. Using Homogenous Time-Resolved Fluorescence (HTRF) and CDK9-cyclin T1 (**Figure 4a**), and Fluorescence Polarisation (FP) (**Figure 4b**) and Isothermal Titration Calorimetry (ITC) (**Figure 4c**) and CDK9-cyclin T2 as orthogonal approaches we confirmed direct binding between CDK9-cyclin T complexes and the BRD4 P-TEFb interacting domain fragment [47] [41] encoding the 52 C-terminal residues of BRD4_1310-1362_. In the HTRF assay format BRD4_1310-1362_ bound to CDK9-cyclin T1 with a measured dissociation constant (K_d_) of 54.6 nM ± 8 nM. In the FP format CDK9-cyclin T2 bound to BRD4 with a Kd of 67.2 ± 4.3 nM. ITC results confirmed the formation of a 1:1 complex between CDK9-cyclin T2 and BRD4 from which a K_d_ of 262 ± 22 nM was determined (**Table 1**).

**Figure 4.**
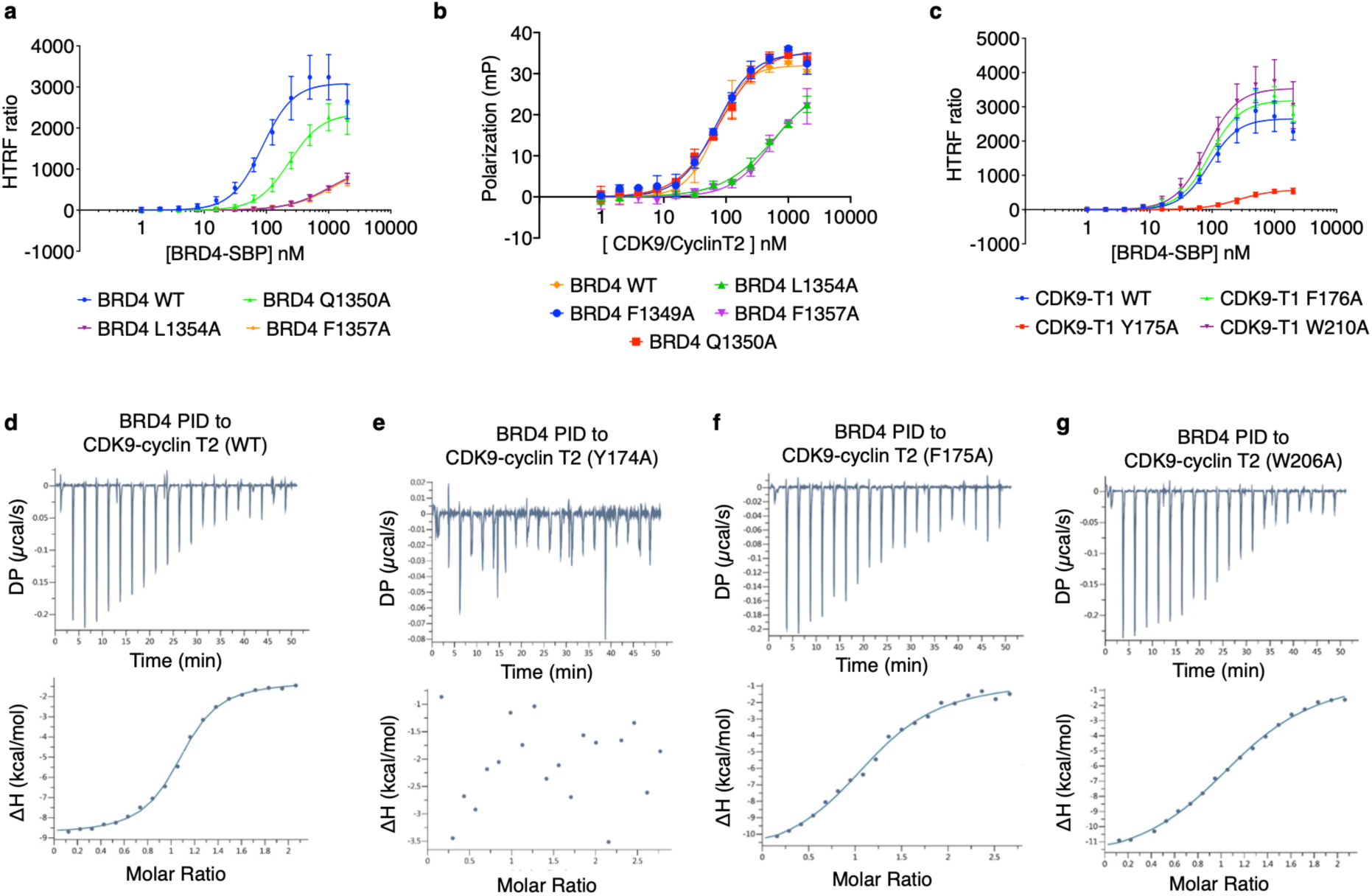
Identification of a cyclin T hotspot required for P-TEFb interaction with BRD4. (**a**) Characterisation of BRD4 mutants that disrupt binding to CDK9-cyclin T1. Mutation of residues Gln1350, Leu1354 and Phe1357, highlighted by the FragLite map and AlphaFold3 model disrupt BRD4 binding to CDK9-Cyclin T1. (**b**) Mutation of BRD4 residues Leu1354 and Phe1357, leads to a loss of BRD4 binding to CDK9-cyclin T2 in a fluorescence polarisation (FP) assay. (**c**) The cyclin T1 Tyr175Ala mutation reduces the Homogenous Time-Resolved Fluorescence (HTRF) signal, whereas the Trp210Ala mutation, previously identified as important for AFF4 interaction and adjacent to Tyr175, shows signals comparable to wild-type. (**d-g**) Isothermal Titration Calorimetry (ITC) plots of CDK9-cyclin T2 complexes with BRD4. Representative titration plots for (d) CDK9-cyclin T2 (e) CDK9-cyclin T2 Tyr174Ala, (f) CDK9-cyclin T2 Phe175Ala (negative control), and (g) CDK9-cyclin T2 Trp206Ala vs the BRD4 P-TEFb Interaction Domain (PID). The top panels show the raw heat signal, and the bottom panels show the integrated heat per injection fitted to a single-site binding model. HTRF experiments were carried out in triplicate and repeated on three separate days. The error bars indicate SD. FP experiments were carried out in triplicate and repeated on three separate days. ITC thermodynamic parameters (K_d_) were derived from three independent biological replicates. Derived K_d_ values are compiled in Table 1. Related to Table 1 and Supplementary Figure 11.

**Table 1.** BRD4 binding to CDK9-cyclin T1 and CDK9-cyclin T2.

| **BRD4** | **CDK9-cyclin T** | **Technique** | **K<sub>d</sub> (nM)** |
| --- | --- | --- | --- |
| BRD4-SBP (1310-1362) WT | CDK9(1-330)/FlagT1(1-281) | HTRF<sup>1</sup> | 86 ± 7 |
| BRD4-SBP (1310-1362) Gln1350Ala | CDK9(1-330)/FlagT1(1-281) | HTRF<sup>1</sup> | 235 ± 16 |
| BRD4-SBP (1310-1362) Leu1354Ala | CDK9(1-330)/FlagT1(1-281) | HTRF<sup>1</sup> | 1030 ± 200 |
| BRD4-SBP (1310-1362) Phe1357Ala | CDK9(1-330)/FlagT1(1-281) | HTRF<sup>1</sup> | 1260 ± 430 |
| BRD4(1310-1362)GFP WT | CDK9 (1-330)/cyclin T2 (1-273) | FP<sup>3</sup> | 67 ± 4 |
| BRD4(1310-1362)GFP Phe1349Ala | CDK9 (1-330)/cyclin T2 (1-273) | FP<sup>3</sup> | 70 ± 6 |
| BRD4(1310-1362)GFP Gln1350Ala | CDK9 (1-330)/cyclin T2 (1-273) | FP<sup>3</sup> | 77 ± 6 |
| BRD4(1310-1362)GFP Leu1354Ala | CDK9 (1-330)/cyclin T2 (1-273) | FP<sup>3</sup> | 650 ± 180 |
| BRD4(1310-1362)GFP Phe1357Ala | CDK9 (1-330)/cyclin T2 (1-273) | FP<sup>3</sup> | 531 ± 130 |
| BRD4(1310-1362)SBP | CDK9 (1-330)/FlagT1 (1-281) WT | HTRF<sup>1</sup> | 90 ± 7 |
| BRD4(1310-1362)SBP | CDK9 (1-330)/FlagT1 (1-281) Tyr175Ala | HTRF<sup>1</sup> | 264 ± 24 |
| BRD4(1310-1362)SBP | CDK9 (1-330)/FlagT1 (1-281) Phe176Ala | HTRF<sup>1</sup> | 92 ± 4 |
| BRD4(1310-1362)SBP | CDK9 (1-330)/FlagT1 (1-281) Trp210Ala | HTRF<sup>1</sup> | 80 ± 6 |
| BRD4 (1310-1362) WT | CDK9 (1-330)/cyclin T2 (1-273) | ITC<sup>2</sup> | 320 ± 90 nM;<br>Molar Ratio= 1.14 ± 0.12<br>(mean ± SD, n = 3) |
| BRD4 (1310-1362) Tyr174Ala | CDK9 (1-330)/cyclin T2 (1-273) | ITC<sup>2</sup> | ND;<br>Molar Ratio: ND |
| BRD4 (1310-1362) Phe175Ala | CDK9 (1-330)/cyclin T2 (1-273) | ITC<sup>2</sup> | 1270 ± 10 nM;<br>Molar Ratio = 1.12 ± 0.10 (mean ± SD, n = 3) |
| BRD4 (1310-1362) Trp206Ala | CDK9 (1-330)/cyclin T2 (1-273) | ITC<sup>2</sup> | 1550 ± 21 nM;<br>Molar Ratio = 1.20 ± 0.06 (mean ± SD, n = 3) |
| BRD4(1310-1362)-SBP | CDK9(1-330)/FlagT1(1-281) | HTRF<sup>1</sup> | 55 ± 8 |
| BRD4(1320-1362)-SBP | CDK9(1-330)/FlagT1(1-281) | HTRF<sup>1</sup> | 82 ± 11 |
| BRD4(1325-1362)-SBP | CDK9(1-330)/FlagT1(1-281) | HTRF<sup>1</sup> | 116 ± 13 |
| BRD4(1331-1362)-SBP | CDK9(1-330)/FlagT1(1-281) | HTRF<sup>1</sup> | 868 ± 150 |
| BRD4(1341-1362)-SBP | CDK9(1-330)/FlagT1(1-281) | HTRF<sup>1</sup> | ND |
| SBP-BRD4(1310-1362) | CDK9(1-330)/FlagT1(1-281) | HTRF<sup>1</sup> | 189 ± 22 |
| SBP-BRD4(1310-1360) | CDK9(1-330)/FlagT1(1-281) | HTRF<sup>1</sup> | 400 ± 60 |
| SBP-BRD4(1310-1357) | CDK9(1-330)/FlagT1(1-281) | HTRF<sup>1</sup> | ND |
| SBP-BRD4(1310-1355) | CDK9(1-330)/FlagT1(1-281) | HTRF<sup>1</sup> | ND |
| FLAG-BRD4(1310-1362) | CDK9(1-330)/SBP-T1(1-281) | HTRF<sup>1</sup> | 26.2 ± 2.9 |
| FLAG-BRD4 (1310-1360) | CDK9(1-330)/SBP-T1(1-281) | HTRF<sup>1</sup> | 52.4 ± 3.8 |
| FLAG-BRD4 (1310-1357) | CDK9(1-330)/SBP-T1(1-281) | HTRF<sup>1</sup> | ND |
| FLAG-BRD4 (1310-1355) | CDK9(1-330)/SBP-T1(1-281) | HTRF<sup>1</sup> | ND |

<sup>1</sup> Homogenous Time-Resolved Fluorescence
<sup>2</sup> Isothermal Titration Calorimetry
<sup>3</sup> Fluorescence Polarisation
ND: K<sub>d</sub> measurement not determined due to poor fit quality or insufficient signal to reliably model binding.

The AF3 models predict that the interaction of BRD4 with cyclin T1 or T2 is centred on BRD4 residues Met1320-Glu1359 (pLDDT scores >90) (**Supplementary Figure 9**). To identify a minimal BRD4 fragment required for CDK9-cyclin T binding and validate the model, a series of BRD4 truncations were generated and fused to a streptavidin-binding protein (SBP) tag positioned at either the N- or C-terminus. BRD4 fragment binding to CDK9-cyclin T1 was subsequently assayed by HTRF (**Supplementary Figure 11**). Truncation of the BRD4 N-terminus to start at residue Arg1325 resulted in little loss in activity, but further deletion to residue Arg1331 markedly reduced binding, and the interaction was no longer detectable when the construct began at Met1341 (**Supplementary Figure 11a**). This result is consistent with the AF3 model which suggests constructs starting at Arg1325, in which pLDDT scores become increasingly higher (>95), retain all observed interactions with CDK9. However, further N-terminal truncation to start at Arg1331 removes many of the salt-bridges associated with CDK9 binding and truncation to Met1341 abrogates CDK9 binding. Similarly, C-terminal BRD4 truncations abolish binding, attributed to loss of cyclin T binding (**Supplementary Figure 11b**). These observations were corroborated by reciprocal experiments in which the SBP tag was incorporated into cyclin T1 (**Supplementary Figure 11c).**

Guided by the AF3 model and FragLite overlay, we selected three BRD4 residues, Gln1350 and Leu1354 (guided by FL2), and Phe1357 (guided by FL2, FL4, FL18, FL19, FL21, FL22, FL23, FL31 and PLG), that make intimate contacts with cyclin T2 in Site 3. Substitution of these residues with alanine in the BRD4_1310-1362_ fragment revealed that while Gln1350Ala was partially tolerated, mutations Leu1354Ala and Phe1357Ala each reduced binding affinity by over tenfold (**Figure 4a** and **Figure 4b**). Consistent with previous reports, Phe1357 is important for both E2 and P-TEFb recognition, whereas Leu1354 appears to be specific to P-TEFb, particularly cyclin T1 [41] [47].

To generate cyclin T point mutants targeting BRD4 interactions, we focused on FragLite-defined residues at Site 3. We selected cyclin T2 Tyr174, which engages in hydrogen bonding and aromatic π-stacking with FragLites; the adjacent Phe175, positioned on the reverse face of H2’ and contacting the Tat helix; and cyclin T2 Trp206 and Trp209, previously identified as disruptive AFF4 mutants [63] but not predicted to affect BRD4 binding. These mutations were made in the context of CDK9-cyclin T1 (**Figure 4c**).

HTRF measurements revealed that the CDK9-cyclin T1 Tyr175Ala mutant significantly reduced BRD4_1310-1362_ binding, whereas the Phe176Ala and Trp210Ala mutations (the latter serving as a negative control) had little or no effect (**Figure 4c**). These findings were confirmed by ITC using equivalent CDK9-cyclin T2 mutants (Tyr174Ala, Phe175Ala, and Trp206Ala) (**Figure 4d-g**). While cyclin T2 Phe175Ala and Trp206Ala retained measurable affinity, binding to the cyclin T2 Tyr174Ala mutant could not be detected, underscoring both site functionality and cyclin T2 Tyr174 as a critical residue for BRD4 engagement.

Collectively, these experiments establish cyclin T1 Tyr175 (cyclin T2 Tyr174) as a critical hotspot for BRD4 recognition, anchoring the interaction within the hydrophobic pocket mapped by FragLites at Site 3 on the cyclin T C-CBF. The mutagenesis data confirm that while neighbouring residues such as cyclin T1 Trp207/cyclin T2 Trp206 and cyclin T1 Trp210/cyclin T2 Trp206 contribute modestly to the interaction with AFF4, they do not abrogate BRD4 binding.

## Discussion

This study demonstrates that highly populated FragLite binding sites correspond to functional protein-protein interaction surfaces. The extent of FragLite occupancy effectively delineates sites of functional importance, distinguishing them from opportunistic fragment-accessible surfaces that can bind fragments but are not sites of authentic protein partner binding. By combining FragLite mapping with AlphaFold modelling and targeted mutagenesis, we show that these fragments not only identify potential interfaces but also provide a roadmap for selectively interrogating protein-protein interactions without exhaustive mutational analysis. Our analysis of cyclin T2 shows that the convergence and density of FragLite binding reliably highlight interaction hotspots, exemplified in this study by the identification of cyclin T1 Tyr175/cyclin T2 Tyr174 as a critical determinant of BRD4 recognition.

This analysis highlights that cyclins that regulate transcription, like those that regulate the cell cycle, exploit multiple discrete protein docking sites (hotspots) to permit protein partners to exploit avidity effects to regulate protein binding, and competition between proteins to effect signal integration [68]. In the case of cyclin T, HEXIM1, a component of the P-TEFb inhibitory complex competes with both BRD4 and AFF4, while the binding sites of BRD4 and AFF4, components of complexes that activate P-TEFb are non-overlapping. Cyclin T function is subverted by the HIV virus through Tat binding to promote viral gene transcription. FragLite Sites 4 and 5 overlap with the Tat helical and tail sequence docking sites respectively suggesting they might, like Site 3, identify PPI sites for cyclin T binding to other authentic partners that HIV has subverted and repurposed. Site 6 is also conserved across cyclin T orthologues and is characterised by rich FragLite binding but does not overlap with any structurally solved cyclin T partner binding site. These unexplored hotspots highlight opportunities for further interrogation and suggest that additional, physiologically relevant interaction surfaces remain to be discovered.

Collectively, these results establish FragLites as a powerful platform for uncovering biologically relevant interaction sites, to guide rational mutagenesis and mechanistic dissection, and to provide a starting point for the design of selective chemical probes. In the cyclin T-BRD4 system, this approach provides a structural and functional framework for exploring targeted modulation of P-TEFb and its broader regulatory network.

### Accession Numbers

The coordinates and the structure factors generated in this study have been deposited in the Protein Data Bank (PDB) https://www.rcsb.org/ under the following accession codes. Other data are available from the corresponding author upon reasonable request.

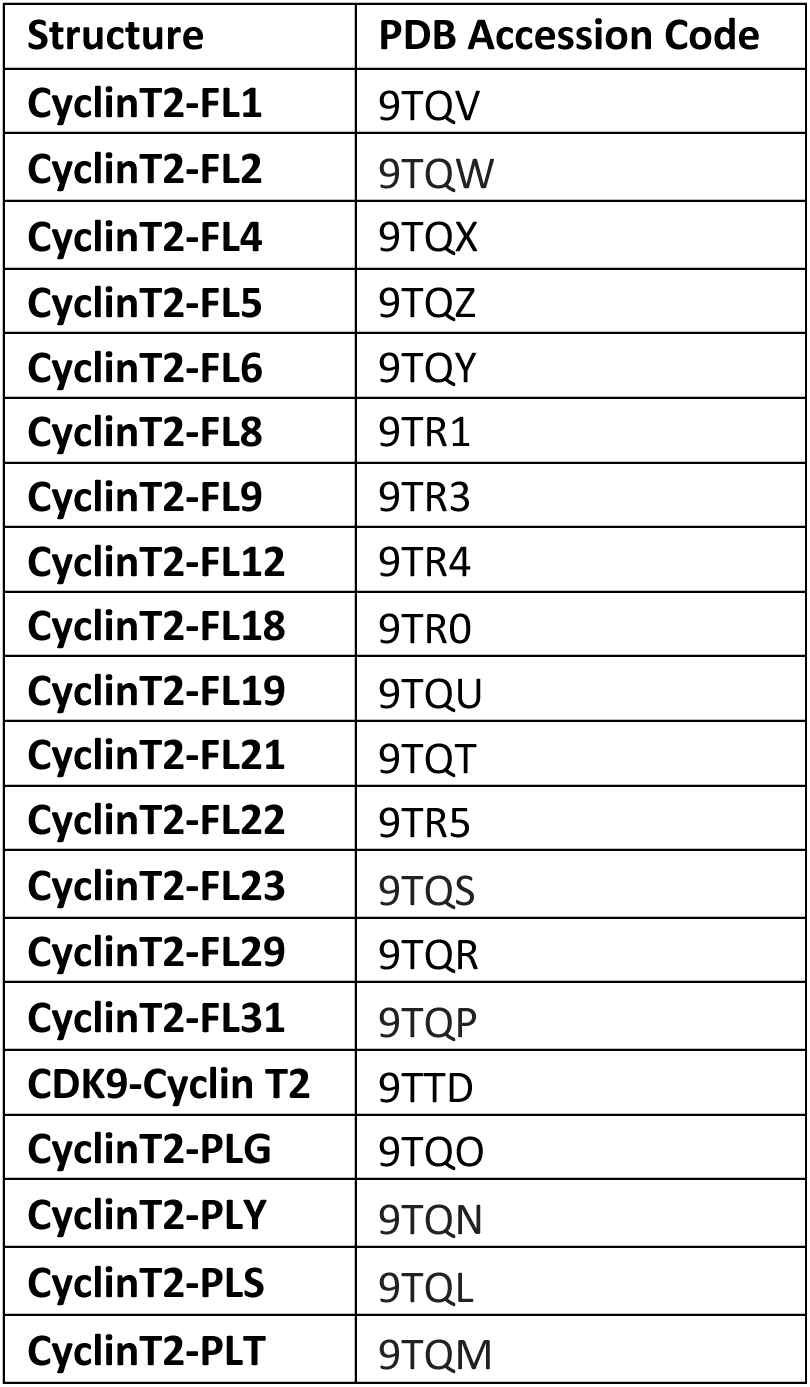

## Acknowledgements

This research was supported by the MRC Grant References G0901526, and MR/N009738/1 (R.H, J.A.E. and M.E.M.N) and Cancer Research UK Grant Reference C2115/A21421. I.H. was supported by a studentship from the Medical Research Council Discovery Medicine North Doctoral Training Partnership. N.J.T.’s position within Newcastle Drug Discovery is supported by Astex Pharmaceuticals. We wish to thank staff on the i04-1 beamline and the XChem Facility (Lab24) at Diamond Light Source (Oxford) for excellent facilities, Dr M Temple for assistance with data collection and Dr S Baumli for providing starting constructs to express CDK9-cyclin T1 and CDK9-cyclin T2. For the purpose of open access, the authors have applied a Creative Commons Attribution (CC BY) license to any Author Accepted Manuscript version arising from this submission.

## Author Contributions

I.H. Conceptualization, methodology, investigation (cloning, protein production, FragLite screening by X-ray crystallography and complex structure determination, writing original draft, review and editing.

R.H. Conceptualization, methodology, investigation (cloning, construct design, protein production, biophysical analysis (HTRF, ITC), writing original draft, review and editing.

N.J.T. Investigation (AlphaFold modelling), review and editing.

A.B. Assistance with structure determination, data collection, Diamond Light Source and with data deposition, review and editing.

M.P.M. Conceptualization, investigation (FragLite design, and fragment screening by crystallography), review and editing.

M.J.W. Conceptualization, investigation (FragLite design), review and editing.

M.E.M.N. Conceptualization, resources, supervision, review, editing and funding acquisition.

J.A.E. Conceptualization, resources, supervision, and funding acquisition, writing original draft, review and editing.

## Declaration of Interests

The other authors declare no competing interests. Some work in the authors’ laboratory is supported by a research grant from Astex Pharmaceuticals.

## Data Access Statement

All crystallographic data and protein structures created during this research are openly available at Protein Data Bank (PDB) https://www.rcsb.org/ through the accession codes provided.

## Materials and Methods

### Construct generation

A codon-optimised DNA sequence encoding the BRD4-CTR (residues 1279-1362) (Uniprot entry O60885) was synthesized by Top Gene Technologies (www.topgenetech.com). PCR fragments of BRD4 (1310-1362), (1320-1362), (1325-1362), (1331-1362) and (1341-1362) were digested with NcoI and SpeI (DNA recognition sequences included in the primer design) and cloned into a modified pET3d vector which contains sequences to express the protein of interest with an N-terminal Glutathione transferase (GST) tag followed by a Human Rhino Virus 3C protease cleavage site (3C), and a C-terminal Streptavidin-Binding Peptide (SBP) tag. PCR fragments of BRD4(1310-1362), BRD4(1310-1360), BRD4(1310-1357), and BRD4(1310-1355) were cloned into modified pET3d vectors that each encode a N-terminal GST tag followed by a 3C protease cleavage site and then either an SBP tag or a FLAG tag preceding the protein of interest. To express CDK9-cyclin T1 and CDK9-cyclin T2 complexes (Uniprot entries CDK9 (P50750), cyclin T1 (O60563) and cyclin T2 (O60583)), a PCR fragment encoding CDK9(1-330) was digested with NcoI and SpeI (DNA recognition sequences included in the primer design) and cloned into a pACEBAC1 vector (Geneva Biotech) modified to include a non-cleavable C-terminal His_6_ tag (CDK9(1-330)His). All mutant proteins were generated using the QuikChange II Site-Directed Mutagenesis Kit (Agilent). The BRD4 mutations Phe1349Ala, Gln1350Ala, Leu1354Ala and Phe1357Ala were introduced into the pet3d vector with BRD4(1310-1362) construct encoding a C-terminal Green Fluorescent Protein (GFP) tag. Cyclin T2 point mutations (Tyr174Ala, Phe175Ala and Trp206Ala) used to carry out the Isotherm Titration Calorimetry (ITC) experiments were introduced into the pNic28-Bsa4 vector encoding cyclin T2 (residues 2-273) preceded by an N-terminal His_6_ tag and TEV protease cleavage sequence (pNic28HisTEVcyclin T2). Cyclin T1 point mutations (Tyr175Ala, Phe176Ala and Trp210Ala) used to carry out the Homogenous Time-Resolved Fluorescence (HTRF) experiments were introduced into a modified pET3d vector encoding an N-terminal His_6_ tag followed by a 3C protease cleavage site preceding the cyclin T1 (1-281) sequence (pET3dHis3Ccyclin T1). All constructs were subsequently verified by sequencing (EuroFins Scientific). pNic28HisTEVcyclin T2 and pNic28HisTEVcyclin T2 (Gln130Arg) used in crystallisation trials were gifts from the Structural Genomics Consortium Oxford.

### Protein expression

GST3cFLAGBRD4, GST3cSBPBRD4 and GST3cBRD4SBP constructs (wild-type and mutants) were expressed in *E. coli* Rosetta DE3 cells (Novagen) in LB broth. Cells were induced at a cell density of OD_600 nm_ *circa* 0.7, with 0.5 mM isopropyl β-D-1-thiogalactopyranoside (IPTG) at 20 °C overnight. *E. coli* cells were then pelleted (4000 xg, 20 min, 4 °C) and re-suspended in 50 mM Tris pH 7.5, 300 mM NaCl, 0.5 mM TCEP supplemented with a protease inhibitor cocktail (Roche) and stored at -20 °C. Cyclin T constructs were expressed in Rosetta2 (DE3) pLysS in LB broth. Cells were grown to an OD_600 nm_ of 0.6-0.8 prior to induction with 0.1 mM IPTG and incubated at 18 °C overnight. Pellets of *E. coli* cells expressing cyclin T1, cyclin T2 or their respective mutants were pelleted (4000 xg, 20 min, 4 °C) resuspended in 50 mM Tris pH 7.5, 300 mM NaCl, 40mM imidazole, 0.5 mM TCEP and protease inhibitors and stored at -20 °C. pACEBAC1CDK9(1-330)His was expressed in accordance with the MultiBac™ system (Geneva-Biotech) methodology. Approximately 5-10 ng of DNA vector was used to transform EmBacY *E. coli* cells harbouring the EMBacY MultiBac™ bacmid (for constitutive expression of yellow fluorescent protein). DNA sequences of interest were transferred to the bacmid via transposition into the mini Tn7 attachment site. White recombinant colonies were selected for subsequent bacmid preparation. MultiBac bacmid DNA was prepared by alkaline lysis (QIAprep Miniprep kit (Qiagen)). The final supernatant was precipitated using isopropanol (40 %) and the resulting pellet was then washed twice with 70 % ethanol, dried and re-suspended in 50 μL of sterile water and 300 μL Insect-Xpress insect cells medium (Lonza) in a sterile hood. 10 μL GeneJuice® Transfection reagent (Merck Millipore) was then added to the DNA and the resulting cocktail was used to infect 0.5 x 10^6^ Sf9 cells per ml (Oxford Expression Technologies Ltd) seeded in 6-well plates. After 48-60 h incubation at 27 °C, the supernatant was collected and positive transfection was verified by fluorescence microscopy. This initial virus stock (Vo) was amplified twice and then used for protein expression. Five ml of V2 virus was added to 1 L of media (Insect-Xpress, Lonza) containing 1 x 10^6^ cells ml^-1^ and then split between 2 x 2.5L glass Erlenmeyer flasks. Cells expressing CDK9(1-330)His were grown at 28 °C, 120 rpm and then harvested by centrifugation (4000 x g, 20 min, 4°C) after 72 h infection. Pellets were re-suspended in 50 mM Tris pH 7.5, 300 mM NaCl, 0.5 mM TCEP and supplemented with a protease inhibitor cocktail (Roche) and stored at -20 °C

### Protein purification

#### CyclinT1 and cyclin T2

Suspended pellets of bacterial cells expressing recombinant cyclin T1 or cyclin T2 were supplemented with 25 µg ml^-1^ lysozyme, 2 µg ml^-1^ DNase and 10 µg ml^-1^ RNase then once thawing was apparent lysed on ice by sonication. After centrifugation at 48,000 xg for 1 h at 4 °C, the clarified lysate was loaded onto a 5 ml HisTrap column (Cytiva) equilibrated in 50 mM Tris pH 7.5, 300 mM NaCl at 4 °C. The column was exhaustively washed with 50 mM Tris pH 7.5, 300 mM NaCl, 40mM Imidazole, 0.5 mM TCEP and bound material was eluted with 50 mM Tris pH 7.5, 300 mM NaCl, 400 mM Imidazole, 0.5 mM TCEP collecting 1 ml fractions. The His tag was removed from cyclin T1 or cyclin T2 by cleaving overnight at 4 °C with either 3C protease (1:100 mass:mass) or TEV protease (1:100 mass:mass) respectively. The cleaved cyclin T samples were further purified by size exclusion chromatography (SEC) (HiLoad 16/60 Superdex 200 pg (Cytiva)) pre-equilibrated in 50 mM Tris pH 7.5, 300 mM NaCl, 0.5 mM TCEP. For cyclin T2, the SEC buffer was supplemented with 5 % ethylene glycol. Fractions containing cyclin T2 or cyclin T1 were identified by SDS-PAGE analysis, concentrated to 10 mg ml^-1^, aliquoted, flash frozen in liquid nitrogen, and stored at –80 °C.

#### CDK9-Cyclin T complexes

Resuspensions of insect cells expressing CDK9(1-330)His were thawed on ice then lysed on ice by sonication. To form the respective CDK9-FLAGcyclinT1 complexes (WT, Tyr175Ala, Phe176Ala and Trp210Ala), and CDK9-cyclinT2 (WT, Tyr174Ala, Phe175Ala, Trp206Ala), the sonicated CDK9 lysate (1 L) was mixed with 10 mg of bacterially expressed cyclin T purified as described above, prior to centrifugation at 40,000 xg for 1 hr at 4 °C. The clarified lysate was filtered and loaded onto a 5 ml HisTrap column (Cytiva) equilibrated in 50 mM Tris pH 7.5, 300 mM NaCl at 4 °C. The column was then exhaustively washed with 50 mM Tris pH 7.5, 300 mM NaCl, 0.5 mM TCEP and bound material was eluted in 1 ml fractions with 50 mM Tris pH 7.5, 300 mM NaCl, 400 mM Imidazole, 0.5 mM TCEP. Fractions containing CDK9-cyclin T complexes were identified by SDS-PAGE analysis, pooled and concentrated prior to further purification by SEC (HiLoad 16/600 Superdex 200 pg (Cytiva) equilibrated in 50 mM Tris pH 7.5, 300 mM NaCl, 0.5 mM TCEP. For crystallisation, SEC fractions containing CDK9-cyclinT2 WT were combined and concentrated to 7 mg ml^-1^, aliquoted, flash frozen in liquid nitrogen, and stored at -80 °C. All other CDK9-cyclin T1/T2 samples were concentrated to 5 mg ml^-1^, aliquoted and flash frozen in liquid nitrogen, and stored at -80 °C. All protein concentrations were determined by NanoDrop2000 UVVis Spectrophotometer (Thermo Scientific) or using a Shimadzu UV1800 spectrophotometer.

#### BRD4

After thawing, *E. coli* cells expressing the various GST3cBRD4 constructs were sonicated on ice in the presence of lysozyme 25 µg ml^-1^, DNase 2 µg ml^-1^ and 10 µg ml^-1^ RNase and centrifuged for 1h at 48,000 g at 4 °C. GST-tagged proteins were purified by applying the filtered supernatant to a 2ml pre-packed GST Sepharose 4B (Cytiva) column at 4 °C. The column was exhaustively washed with 30 ml of lysis buffer (50 mM Tris pH 7.5, 300 mM NaCl, 0.5 mM TCEP). Resin was resuspended in the column in 10 ml of lysis buffer supplemented with 0.05 mg ml^-1^ 3C protease and left overnight at 4 °C. The column was drained and the flowthrough collected. The column was then washed with 10 ml of lysis buffer and the second flow through added to the first. The combined flow-through was concentrated and further purified by SEC (HiLoad 16/60 Superdex 75 pg (Cytiva)). The choice of buffer for SEC purification depended on the application. For subsequent analysis by ITC, the SEC buffer was 20 mM HEPES pH 7.5, 300 mM NaCl, 0.5 mM TCEP whereas for HTRF or FP analysis the SEC column was equilibrated in PBS pH 7.5. Fractions containing BRD4 were concentrated, aliquoted, flash frozen in liquid nitrogen, and stored at –80 °C. Protein concentrations were estimated using Coomassie protein assay reagent (Pierce) calibrated with BSA solution.

### Biophysical assays

#### Homogenous Time-Resolved Fluorescence (HTRF)

Assay proteins and reagents were prepared in PBS pH 7.5, 0.005 % Tween 20. Experiments were carried out over 12 serial dilution points at varying concentrations of BRD4-SBP. CDK9-FLAGcyclin T1 (5 nM) was mixed at a 1:1 volume ratio (5 μl, total volume 10 μl) with a Terbium (Tb)-anti-FLAG antibody (260 ng mL^-1^ (CisBio)) and incubated together for 60 min at 20 °C. BRD4-SBP (8 µM) and streptavidin-tagged XL665 dye (250 nM, 1/32^nd^ BRD4-SBP concentration (CisBio)) were mixed at a 1:1 volume ratio (5 μl) and serial diluted to form a dilution series ranging from 0-2 µM BRD4-SBP. Once incubated, the Tb-labelled anti-FLAG antibody/FLAG-cyclin T complex was again mixed in equal parts with the BRD4-SBP/SAXL665 to form a CDK9-FLAGcyclin T1-BRD4 complex. The plate was incubated standing at 20 °C for an additional 60 min and scanned.

In the reverse format, assay components were prepared as described above, with the exception that CDK9-FLAGcyclin T1 was titrated while BRD4-SBP was held constant. BRD4-SBP (5 nM) was mixed at a 1:1 volume ratio (5 μl, total volume 10 μl) with streptavidin-tagged XL665 dye (250 nM, CisBio) and incubated for 60 min at 20 °C. CDK9-FLAGcyclin T1 (8 µM) was mixed with Terbium (Tb)-anti-FLAG antibody (260 ng mL⁻¹, CisBio) at a 1:1 volume ratio (5 μl) and serially diluted over 12 points to generate a concentration range of 0-2 µM. Following incubation, both solutions were combined in equal volumes to form the CDK9-FLAGcyclin T1-BRD4 complex. The plate was incubated at 20 °C for a further 60 min prior to measurement. Samples were excited using a wavelength of 337 nm and emission spectra measured at 620 nm and 665 nm (PHERAstar FS (BMG LABTECH)).

#### Isothermal Titration Calorimetry

CDK9-cyclin T2 was buffer exchanged using a HiTrap desalting column (5 mL) (Cytiva) into ITC buffer (50 mM HEPES, 300 mM NaCl, 0.5 mM TCEP, pH 7.4) and protein concentration was then determined using a Nanodrop 2000 at an absorbance of 280 nm with sequence derived extinction coefficients (http://web.expasy.org/ protparam/). The buffer exchange was carried out to alleviate subsequent heats of dilution associated with using Tris buffer in the ITC experiments. CDK9-cyclin T2 was prepared in ITC buffer in the cell at 10 μM with BRD4 at 100 uM in ITC buffer in the syringe. All ITC experiments were carried out at 20 °C using a Microcal PEAQ ITC instrument (Malvern) with 1× injection of 0.4 μL for 0.8 s followed by 19× 2 μL, 4 s injection with 120 s spacing between injections, using a reference power of 6 μcal s^-1^. Data were analysed using MicroCal PEAQ Evaluation software (Malvern) using the one set of sites model to calculate the dissociation constant and the error of the fit. Experiments were carried out in triplicate with three separate biological preparations of CDK9-cyclin T2.

#### Fluorescence Polarisation

BRD4-GFP (10 nM) was incubated with a 12-fold serial dilution of CDK9-cyclin T2 (0-2 μM). Reagents were prepared in PBS with 0.005 % Tween 20. The plate was incubated at room temperature for 30 min before being scanned. Samples were excited using a wavelength of 480 nm and emission spectra measured at 520 nm (PHERAstar FS (BMG LABTECH)).

Binding curves were plotted using GraphPad Prism 6 from which the dissociation constants( K_d_s) were determined. The curves shown are representative binding curves from at least two runs with each mutant carried out on separate days.

### Protein crystallisation, FragLite mapping and structure determination

CyclinT2 (2-273 Gln130Arg) was concentrated to 9-10 mg ml^-1^ prior to setting up crystallisation plates. Initial crystallization conditions were identified in the Morpheus sparse matrix screen following screening against a wide range of commercially available screens using the sitting drop vapour diffusion method in 96-well 3-drop SWISSCI plates containing 50 µl precipitant buffer. Typically, purified cyclin T2 protein was mixed at a ratio of 1:1, 1:2 and 2:1 with precipitant solution to a total volume of 600 nL using a Mosquito robot (SPT Labtech) and incubated at 20 °C. Single crystals were grown from a matrix containing 0-14 % PEG 10,000, 18-28 % ethylene glycol, 0.12 M Morpheus ethylene glycol mix, 0.1 M Morpheus buffer system 3 pH 8.5 at 20 °C for a minimum of 5 days. Crystals typically grew as plates of 100-300 μm x 30-60 μm and so matched the focused beam size (60 x 50 μm).

Soaking of the FragLite Library (CancerTools.org) into single crystals was conducted at the XChem Facility, Diamond Light Source ((DLS), Oxfordshire, UK). Crystals were soaked for 2 h in 50 mM compound prepared by ECHO dispensing 60 nL stock solution (500 mM compound in DMSO) into the crystal drop. Crystals were harvested directly from the drop and flash cooled in liquid nitrogen. All data sets were collected by automated data collection on beamline I04-1 with X-ray centring using a strategy comprising 360 ° rotation, 0.24 ° oscillation and 0.04 s exposure.

A starting model was obtained from PDB code 2IVX (cyclin T2 [48]). Images were processed automatically at DLS using AutoProc [87] and Xia2 with DIALS [88] within the XChem Explorer pipelines [75,76] and hits identified using the Pan-Dataset Density Analysis (PanDDA) graphical user interface [77]. FragLite datasets were further processed locally using the CCP4i2 graphical user interface [89]. Ligands were prepared using AceDRG [90]. Log-likelihood gain (LLG) anomalous signal maps were derived from molecular replacement using PHASER [91]. Phases were further refined through iterative rounds of model building in COOT [92] refinement in REFMAC5 [93] and validated using Molprobity [94]. Ligand-bound structures and their associated structure factors have been deposited in the PDB with accession codes.

Prior to crystallisation, CDK9(1-330)His-cyclin T2(1-273) protein aliquots were supplemented with 20% ethylene glycol v/v. Crystals of CDK9-cyclin T2 were identified in the PACT premier^TM^ sparse matrix screen (Molecular Dimensions) following screening against a wide range of commercially available screens using the sitting drop vapour diffusion method in 96-well 2-drop SWISSCI plates containing 80 µl precipitant buffer. Protein was mixed in a 1:1 and 1:2 ratio with precipitant buffer to a total volume of 200 nL using a Mosquito robot (SPT Labtech) and incubated at 20 °C. Single crystals were grown as rods from a matrix containing 6-9 % PEG 3350, 0.2 M sodium acetate pH 7.0 at 20 °C for a minimum of 7 days. X-ray diffraction data were recorded using Beamline i-03 at Diamond Light Source, Oxford, UK. Dataset was automatically processed using DIALS [88] and analysed locally using the CCP4i2 graphical user interface [89]. A starting model of CDK9-cyclinT1 was obtained from the PDB (PDB 3IMY [48]). Data were processed using DIALS while PHASER [91] was employed for phasing and REFMAC [93] used during refinement. Model building was performed using COOT [92]. Figures were prepared using CCP4MG [95].

## AlphaFold Structure Prediction

Structural models of CDK9-Cyclin T1-BRD4 and CDK9-Cyclin T2-BRD4 were generated by using the AlphaFold3 server [96]. queried with the full-length sequence for CDK9 (P50750), cyclin T1 (O60563, 1-281) or T2 (O60583, 1-273), and the BRD4 PID (O60885, 1309-1362). For HEXIM1, models were generated using two copies of the C-terminal region of HEXIM1 predicted to form a coiled-coil structure as a dimer (O94992, 255-359). Results were visualised in ChimeraX [97] and further analysed using AlphaBridge [98].

## Supplemental Information

Supplementary Figures S1-S11, Supplementary Tables S1-S5.

Details of all resources used or generated in this study are provided in Supplementary Table S5.

## Declaration of generative AI and AI-assisted technologies in the manuscript preparation process

During the preparation of this work, the authors used Microsoft Copilot in ideation and some instances of support in simplifying technical language. After using this tool/service, the authors reviewed and edited the content as needed and take full responsibility for the content of the published article.

## Abbreviations

CBD: cyclin box domain
CBF: cyclin box fold
CDK: cyclin-dependent kinase
CTD: RNA polymerase II C-terminal domain
DSIF: dichlorobenzimidazole riboside-sensitivity-inducing factor
Fluorescence Polarisation: (FP)
HIV: human immunodeficiency virus
Homogenous Time-Resolved Fluorescence: (HTRF)
IDR: Intrinsically Disordered Region
Isothermal Titration Calorimetry: (ITC)
NELF: negative elongation factor
PanDDA: Pan-Dataset Density Analysis
PPI: protein-protein interactions
P-TEFb: positive transcription elongation factor b
PID: P-TEFb-interacting domain
RNAPII: RNA polymerase II
TAT: Trans-Activator of Transcription
TRM: Tat-TAR recognition motif.

## Notes

### Competing Interest Statement

The authors declare no competing interests. Some work in the authors laboratory is supported by a research grant from Astex Pharmaceuticals.

