## Supplemental Material for "FragLite mapping to identify the BRD4 recruitment site of P-TEFb"

| |
|---|
| Figure S1: Sequence alignment of cyclin T1 and cyclin T2 isoforms |
| Figure S2: Structural comparison of P-TEFb complexes |
| Figure S3: Data collection and refinement statistics |
| Figure S4: The PepLite map of Cyclin T2 |
| Figure S5: Exemplar modelling of FragLites in electron density maps |
| Figure S6: FragLite and PepLite binding to cyclin T2. |
| Figure S7: Protein interactions at the N-terminal cyclin box fold H1 helix |
| Figure S8: FragLite binding to cyclin T2 at the CDK9 interface. |
| Figure S9: The Alphafold model of the CDK9-cyclin T-BRD4 complex |
| Figure S10: The HEXIM1 binding site on cyclin T |
| Figure S11: Biophysical characterisation of the interaction between CDK9-cyclin T complexes and BRD4 |
| Table S1: FragLite/PepLite structures, IUPAC names and SMILES string. |
| Table S2: Data statistics and refinement details |
| Table S3: Summary of FragLite binding events categorised by detection method and associated anomalous signal RMSD values |
| Table S4: Summary of FragLite binding events at partner protein interaction sites |
| Table S5: Key resources |

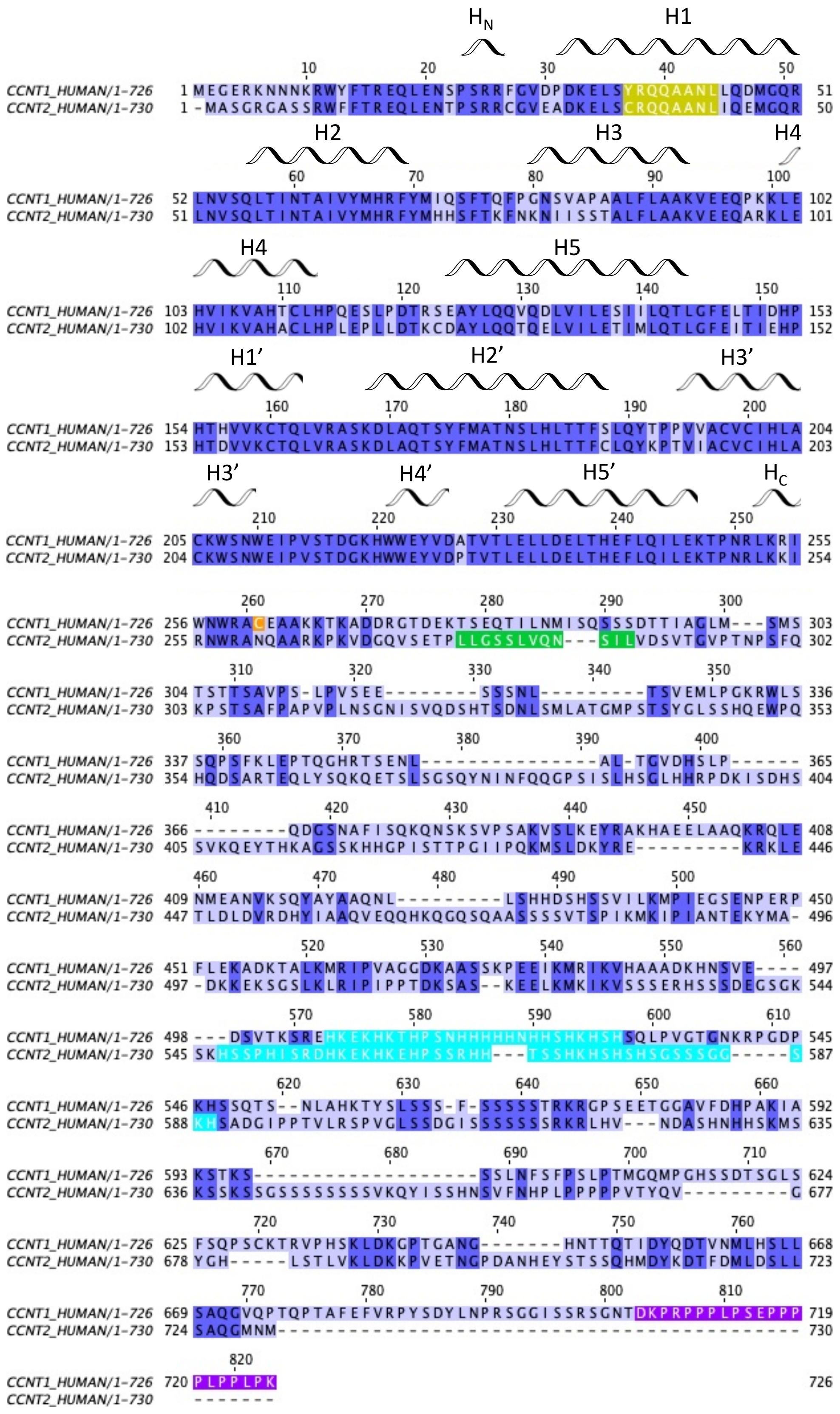

**Supplementary Figure 1. Sequence alignment of cyclin T1 and cyclin T2 isoforms.** Sequence alignment of human cyclin T1 (UniProt: O60563) and human cyclin T2 (UniProt: O60583) carried out using Clustal Omega [1]. Sequences were processed and coloured in Jalview [2] using the % identity function to illustrate isoform sequence conservation. Additional colours highlight key sequence features: MRAIL equivalent sequence (yellow), Cys261 (orange), Leu-rich stretch (green), His-rich stretch (cyan), PEST sequence (purple). Related to Figure 1.

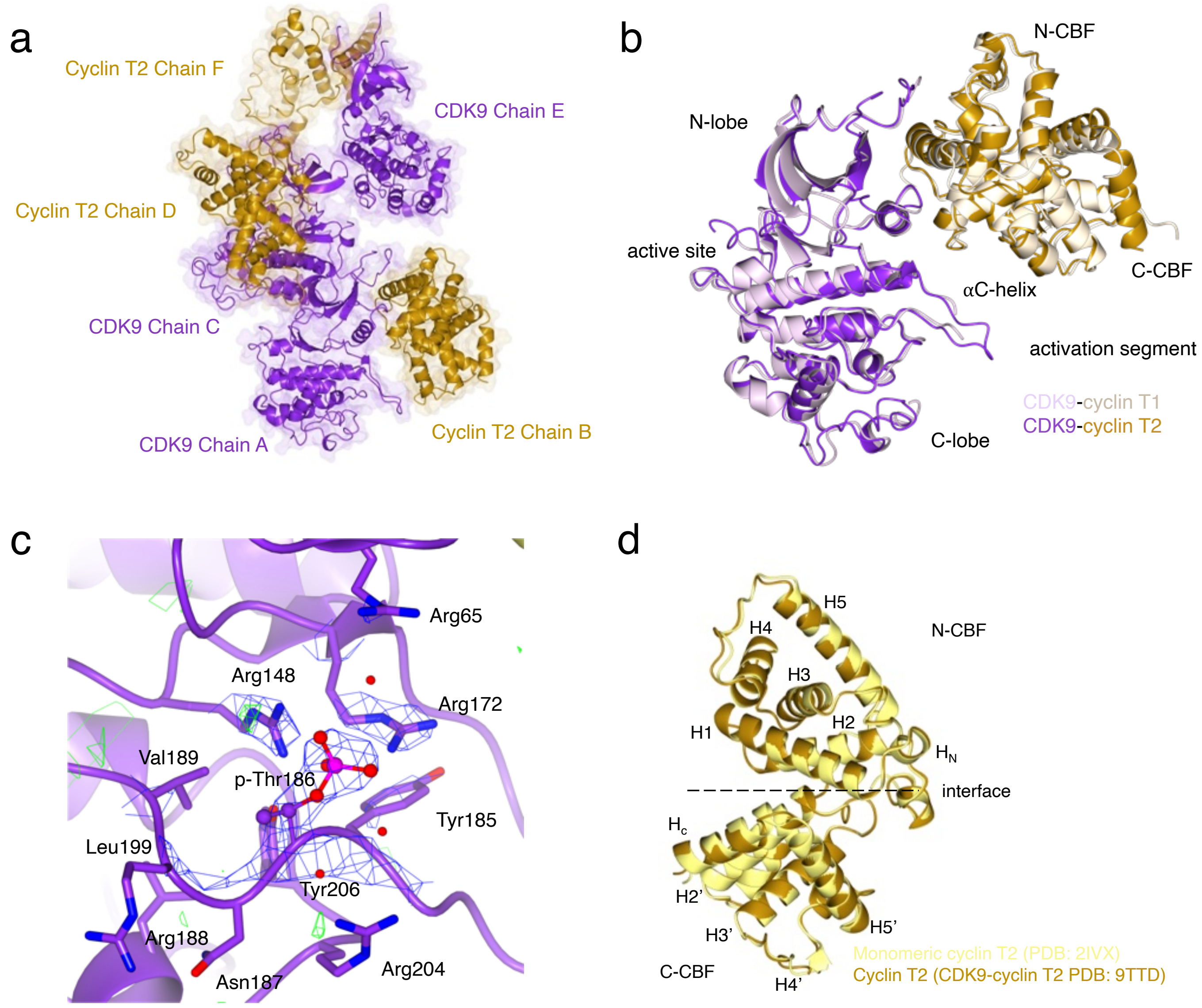

**Supplementary Figure 2. Structural comparison of P-TEFb complexes. (a)** CDK9-Cyclin T2 crystallises with three copies of the dimer in the asymmetric unit. CDK9 and cyclin T2 are shown as purple and goldenrod ribbons, respectively, with a faded molecular surface**. (b)** An overlay of the crystal structure of CDK9-cyclin T1 (pale purple and cream ribbons, PDB 4IMY) with the crystal structure of CDK9-cyclin T2 (PDB 9TTD) shows conservation of the mode of binding of cyclin T isoforms to CDK9 and of CDK activation. Interactions are localised to the protein N-lobes and CDK9-cyclin T2 exhibits an identical splayed disposition of the two subunits. **(c)** The 2Fo-Fc electron density map shows strong electron density for phosphorylated Thr186 (p-Thr186) in the activation segment which engages through electrostatic interactions with Arg148 and Arg172 and with Arg65 and Arg204 through water-mediated hydrogen bonds. The weighted 2Fo-Fc map contoured to 0.79 e/Å$^3$, 2.8 rmsd and the Fo-Fc difference map ± 0.31 e/Å$^3$, 1.1 rmsd. **(d)** A comparison of the monomeric cyclin T2 structure (yellow ribbon, PDB 2IVX) with the cyclin T2 subunit when bound to CDK9 (goldenrod ribbon) shows no significant change in the cyclin T2 conformation (rmsd 0.60 Å over 255 Ca atoms). Figure prepared using CCP4mg [3]. Related to Figure 1.

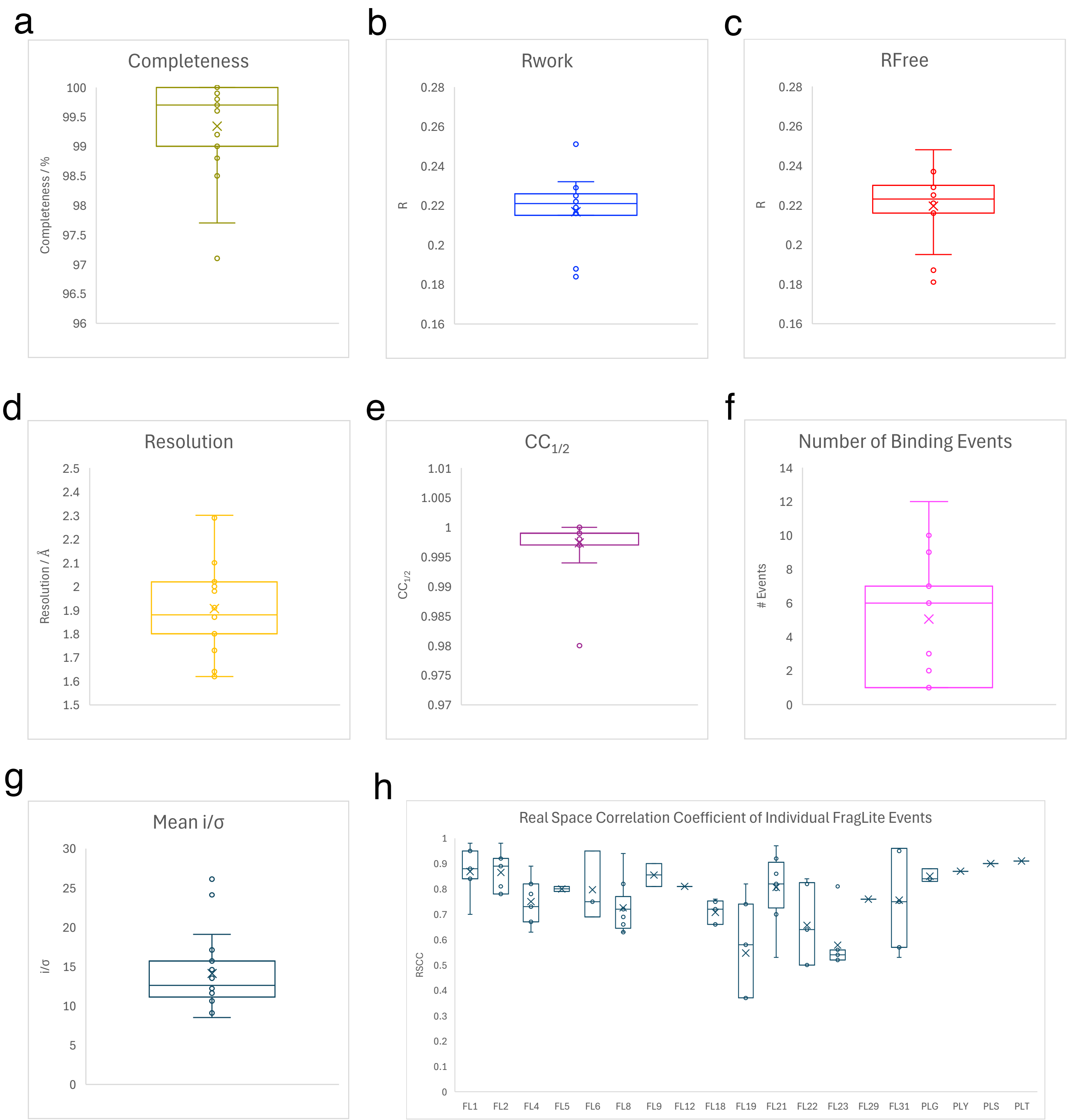

**Supplementary Figure 3. Data collection and refinement statistics. (a)** Per-structure statistics presented as box plots reflecting mean and range. (**b)** Real-space correlation coefficients (RSCC) for each FragLite binding event compiled to illustrate mean and SD. Related to Figure 1.

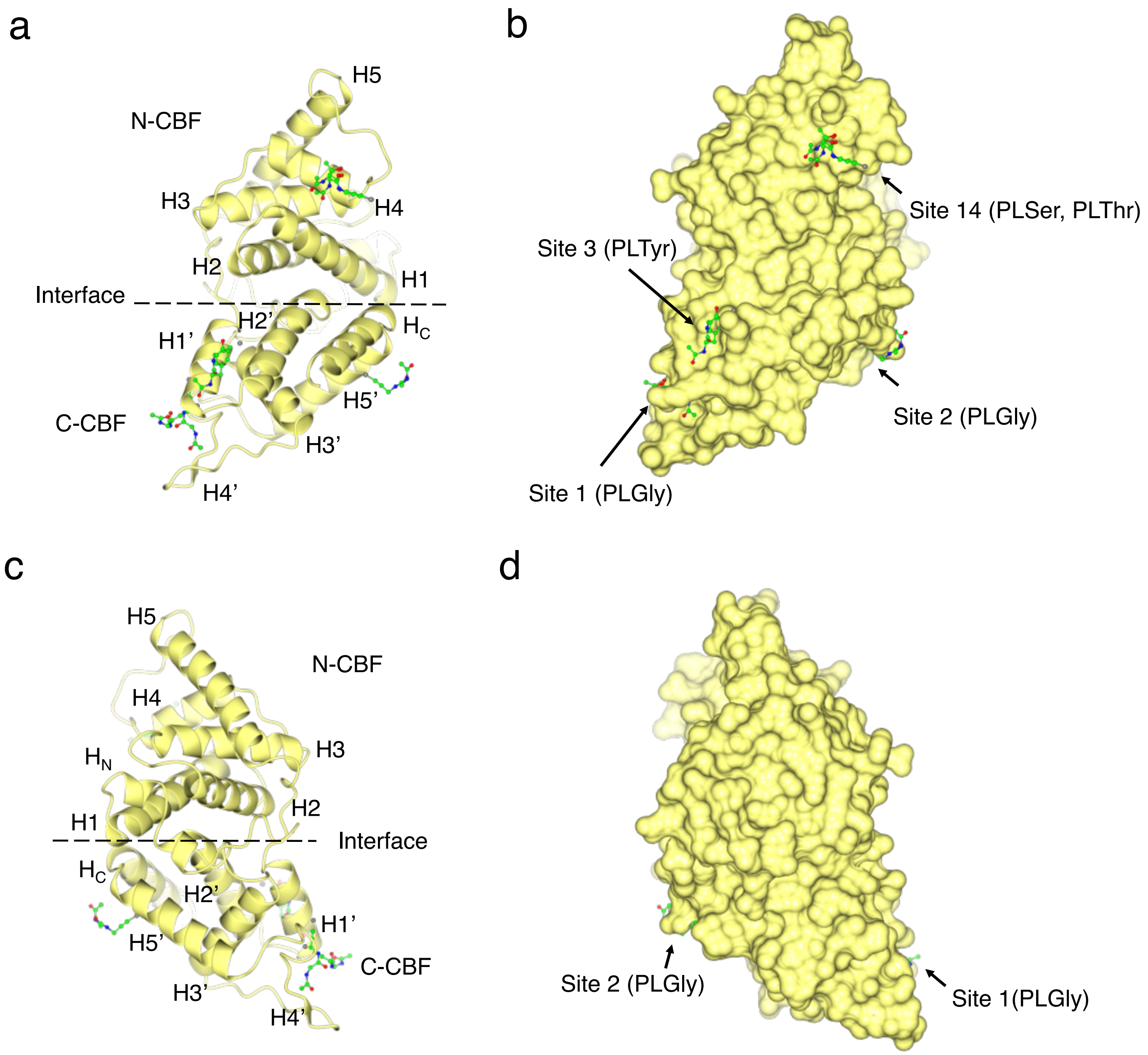

**Supplementary Figure 4. The PepLite map of cyclin T2. (a, c)** PepLite binding events on monomeric cyclin T2 (lemon ribbon) overlap FragLite-identified hotspot sites, showing preferential engagement of the C-terminal cyclin box fold (C-CBF). **(b, d)** Molecular surface of cyclin T2 showing site-specific PepLite occupancy. PepLiteGly binds the two most populated FragLite sites (Site 1, 2 events and Site 2, 1 event). PLTyr occupies Site 3, and PLSer/PLThr define a novel site on the N-terminal cyclin box fold (N-CBF) (Site 14). PepLite binding enriches the FragLite map by providing additional chemical diversity at the most highly populated sites. Figure prepared with CCP4mg [3]. Related to Figure 1.

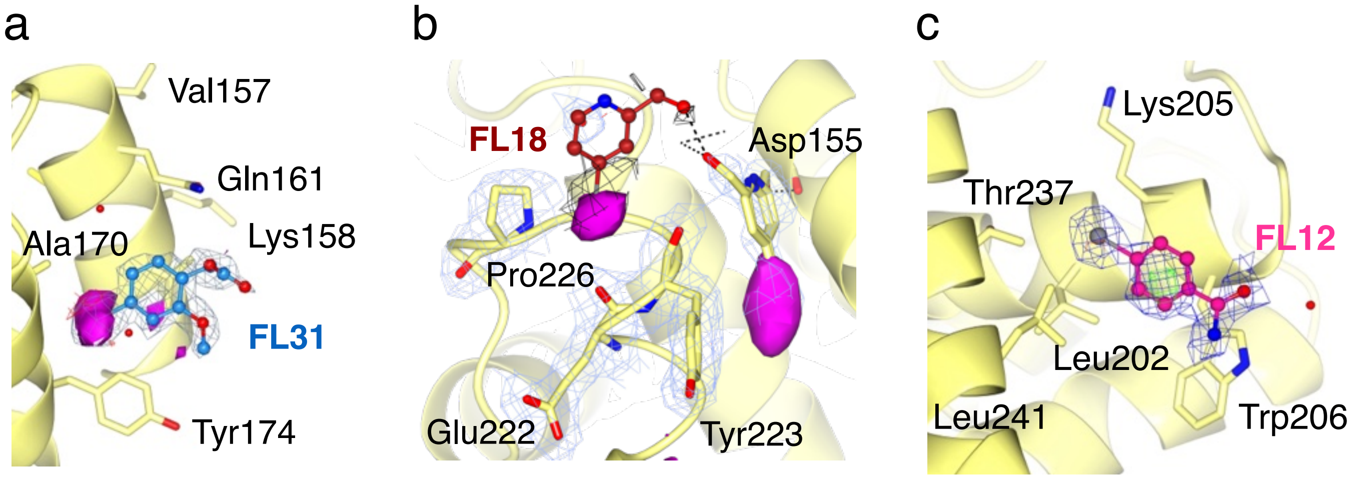

**Supplementary Figure 5. Exemplar modelling of FragLites in electron density maps.**
**(a)** An example of a PanDDA event map (grey mesh) equipped with the anomalous electron density map (magenta sphere) shows unambiguous binding of FL31 (dodger blue). The weighted 2Fo-Fc map contoured to 0.6 e/Å$^3$, 1.1 rmsd and the Fo-Fc difference map ± 0.54 e/Å$^3$, 1 rmsd. **(b)** An example of ambiguous FragLite binding derived from a PanDDA event map for FL18 (shown in maroon and electron density as black mesh). The addition of the anomalous electron density map (magenta sphere) supports the binding of FL18 in this pocket. Electron density contoured to 0.3 e/Å$^3$, 0.6 rmsd. **(c)** The PanDDA event map details binding of FL12 (pink) when anomalous density is absent. Electron density contoured to 0.45 e/Å$^3$, 0.9 rmsd. Figure prepared with CCP4mg [3]. Related to Figure 1.

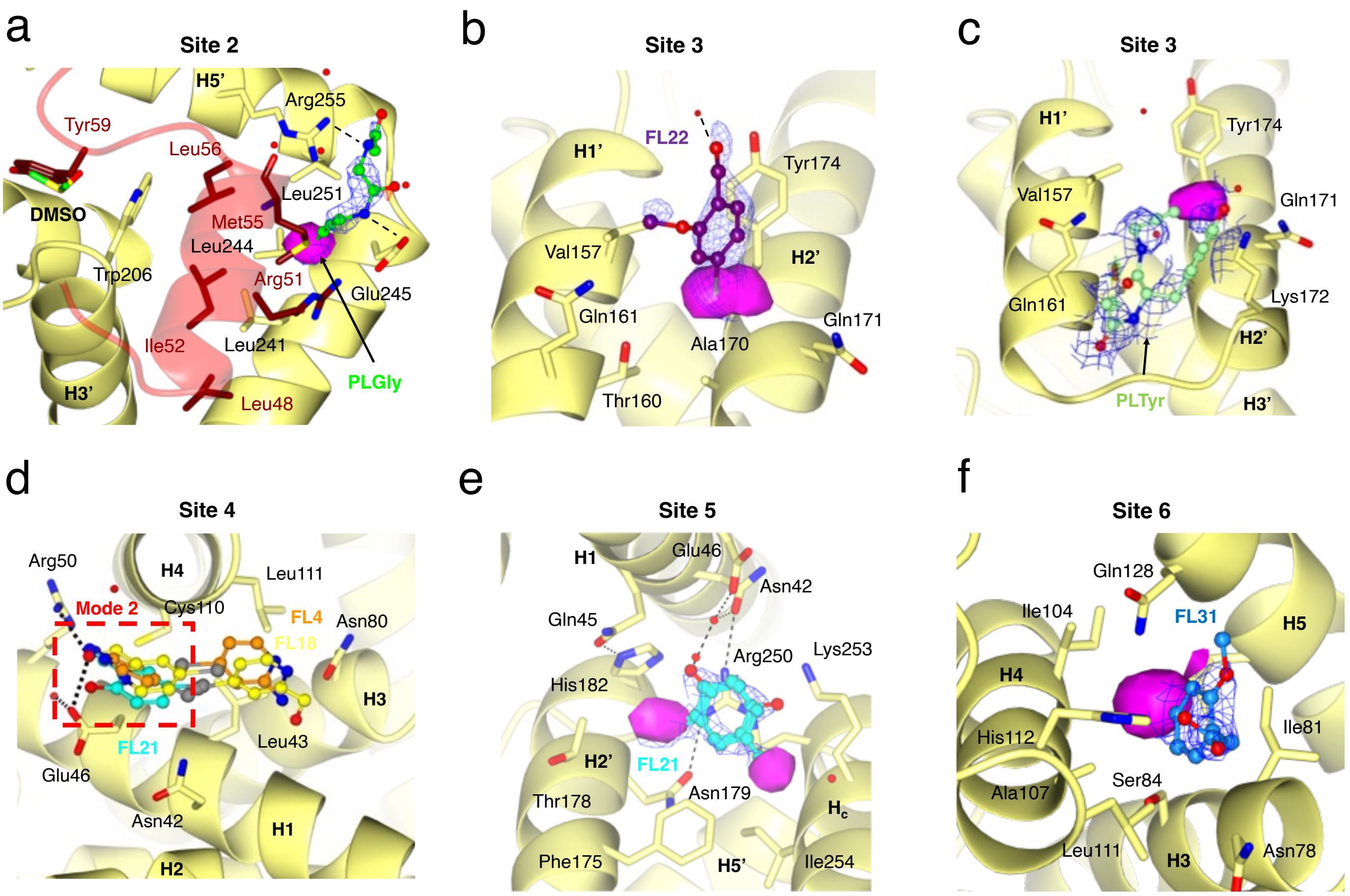

**Supplementary Figure 6. FragLite and PepLite binding to cyclin T2. (a)** PepLiteGly (PLGly) forms hydrogen bonds with cyclin T2 Arg255 and Glu245, mimicking contacts made by AFF4 residues Arg51 and Met55. Electron density shown is the weighted 2Fo-Fc map contoured to 0.41 e/Å$^3$, 1.48 rmsd and Fo-Fc difference map at ±0.29 e/Å$^3$ (1.04 rmsd). **(b)** FragLite occupancy at the Site 3, the "Tat Loop" site. FragLites occupy a site near cyclin T2 Tyr174, burying bromine substituents in a hydrophobic pocket that mimics the Tat Pro3 side chain. For FL22, binding is reinforced by a displaced π-π stack with Tyr174 and an H-bonding network involving Val157 and adjacent water molecules. The weighted 2Fo-Fc map is contoured to 0.23 e/Å$^3$, 0.86 rmsd. **(c)** PLTyr binding at cyclin T2 Site 3, the "Tat Loop". The PepLite bromine atom overlaps spatially with the bromine burial site of neighbouring FragLites, as observed in the overlay map. Additional stabilising interactions span the pocket including an acetamide-carbonyl hydrogen bond with Gln161, and a cation-π interaction between the PepLite phenyl ring and Lys167. The weighted 2Fo-Fc map contoured to 0.21 e/Å$^3$, 0.7 rmsd and Fo-Fc difference map at ±0.32 e/Å$^3$ (2.93 rmsd). **(d)** Alternative FragLite binding at Site 4, the "Tat Tail" site. FL4 (orange), FL18 (yellow), and FL21 (cyan) adopt a second binding mode. In Binding Mode 2, these compounds utilise hydrogen-bonding substituents to coordinate with polar residues Glu46 and Arg50. **(e)** FragLite binding at Site 5 the "Tat Helix" site is driven by electrostatic and H-bonding interactions. FL21 exhibits a dual overlapping mode where the phenyl core forms a cation-π interaction with Arg250 and, in one orientation, facilitates an H-bond network to Asn42 and Glu46 via a water molecule. Electron density shown is the weighted 2Fo-Fc map contoured to 0.8 e/Å$^3$, 1.42 rmsd. **(f)** FragLite binding at Site 6. This site is located on the N-terminal Cyclin Box Fold (N-CBF) within a deep pocket formed by helices H3, H4 and H5 Binding is supported through hydrophobic burial of the aromatic core and halogen into a pocket consisting of Ile104, Ala107 and Ile81. Hydrogen bonding interactions are also supported between the His112 sidechain and the carboxylate of FL31. The weighted 2Fo-Fc map contoured to 0.40 e/Å$^3$, 1.85 rmsd. Figure prepared using CCP4mg [3].

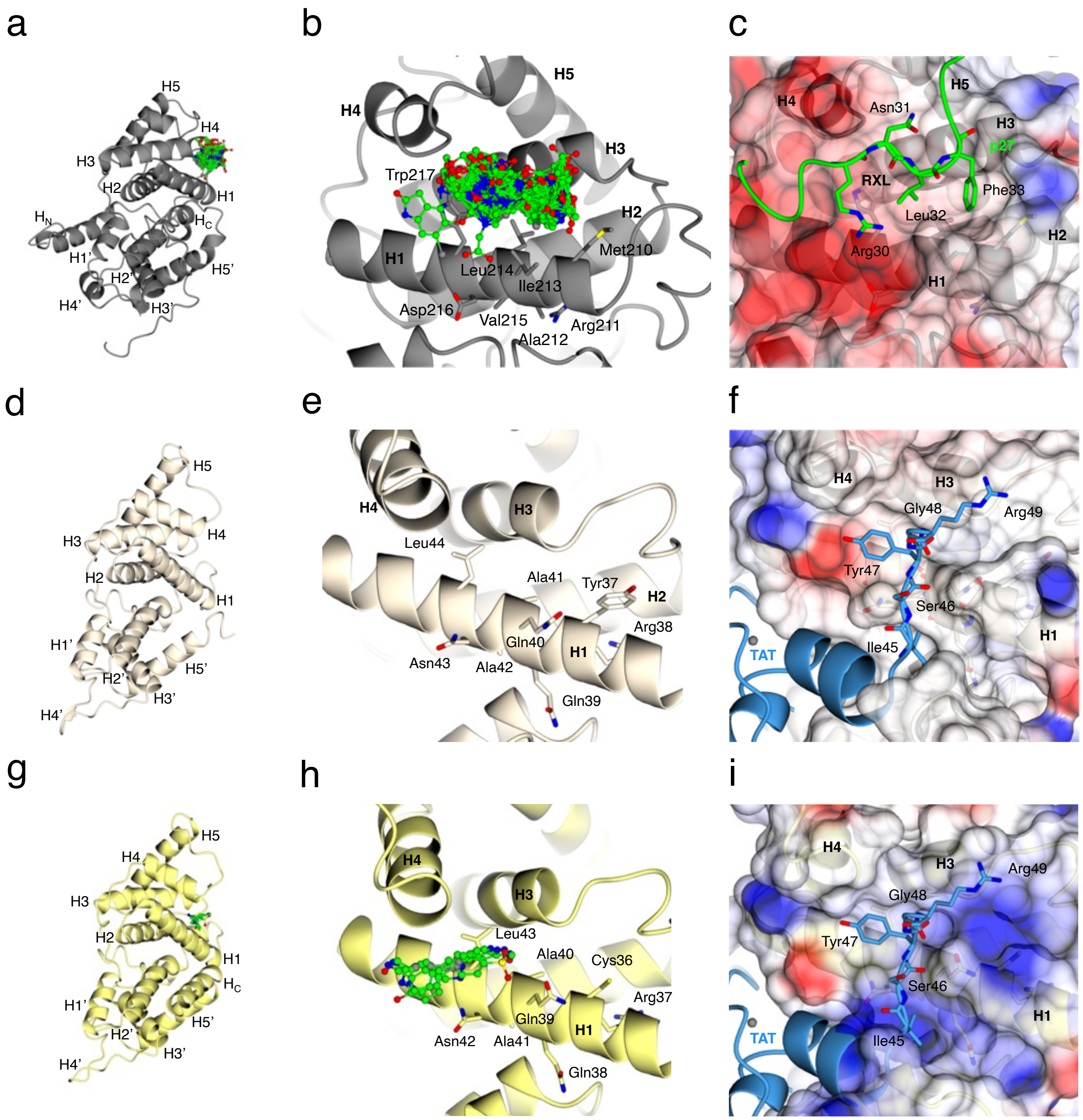

**Supplementary Figure 7. Protein interactions at the N-terminal cyclin box fold H1 helix. (a-c) Cyclin A2.** (a) Structure of cyclin A2 highlighting the H1 helix that contains the MRAIL sequence and contributes to the cyclin A recruitment site that recognises the RXL motif. (b) FragLite map at the cyclin A2 RXL binding site derived from crystallographic fragment screening [4]. (c) Comparison with the binding of p27KIP1 (green) to cyclin A2 (PDB 1JSU). **(d-f) Cyclin T1.** (d) Overall structure of cyclin T1 (beige) with α-helices labelled, highlighting the position of the H1 helix. (e) Close-up of the H1 helix. (f) Electrostatic surface representation of cyclin T1 with the HIV-1 Tat peptide bound (PDB 3MI9 (blue)), illustrating engagement of HIV Tat residues Ile45–Arg49 along the H1 helix. **(g-i) Cyclin T2.** (g) Overall structure of cyclin T2 with the conserved H1 helix highlighted. (h) Zoomed view of the authentic FragLite map at the H1 helix as determined in this study. (i) Electrostatic surface of cyclin T2 overlaid with the Tat peptide (blue) positioned from the cyclin T1–Tat complex (PDB 3MI9). Panels (a), (d) and (g) are in the same orientation and related to the other panels by a rotation 90 ° about the y-axis. Figure prepared using CCP4mg [3]. Related to Figure 2.

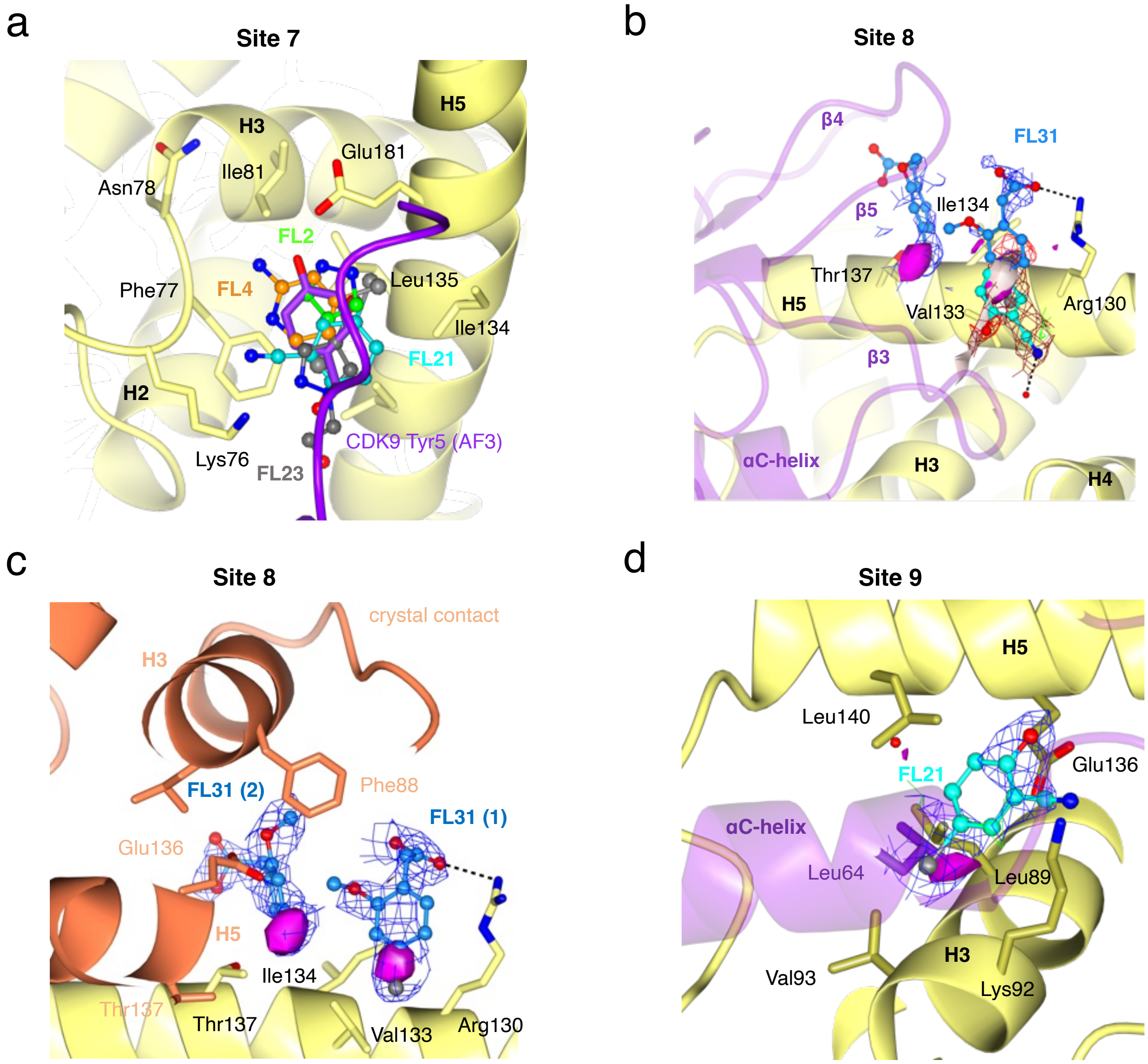

**Supplementary Figure 8. FragLite binding to cyclin T2 at the CDK9 interface.** (**a**) FragLite binding at Site 7, the "CDK9 N-terminus". FL2 (green), FL4 (orange), FL21 (cyan) and FL23 (grey) bind at the H2-H3 linker, H5 helix interface on cyclin T2. FragLite cores overlap with the CDK9 N-terminal Tyr5 position (shown in purple), as observed in the Alphafold3 model [5], suggesting a docking surface for the flexible N-terminal segment. (**b-c**) FragLite binding at Site 8, "CDK9 b sheets". Superposition of Fragment hits FL21 (cyan) and FL31 (dodger blue) reveals three binding events that cluster along the surface shown to bind the b3-b4-b5 sheet region of CDK9 (purple ribbon). FL31 anomalous magenta (blue density at 0.22e/Å$^3$, 1.01 rmsd) FL21 anomalous pale pink, tan density (0.22e/Å$^3$, 0.86 rmsd), difference 0.29e/Å$^3$, 3rmsd. (c) FL31 adopts two distinct orientations at Site 8. In Binding Mode 1, the carboxylate tail forms an ionic interaction with Arg130, while the bromine substituent is buried between Val133 and Arg130. In Binding Mode 2, FL31 is wedged between Ile134/Val137 of the main chain and Phe88/Lys92/Glu136 of the crystal contact (shown in orange). Here, the methoxy substituent overlaps with the Lys92 sidechain (not shown), disrupting a salt bridge with Glu136. (**d**) FragLite binding at Site 9, "CDK9 a-helix". A single fragment (FL21, cyan) interacts with a region supporting stabilisation of the CDK9 aC (H1) helix, with the fragment's bromine atom overlapping Leu64 at the aC helix. Weighted 2Fo-Fc map is contoured at 0.18 e/Å$^3$ (0.7 rmsd) and Fo-Fc difference map at ±0.29 e/Å$^3$ (3 rmsd). Anomalous difference density peak for FragLite detection is represented as a solid magenta sphere. Figure prepared using CCP4mg [3]. Related to Figure 2.

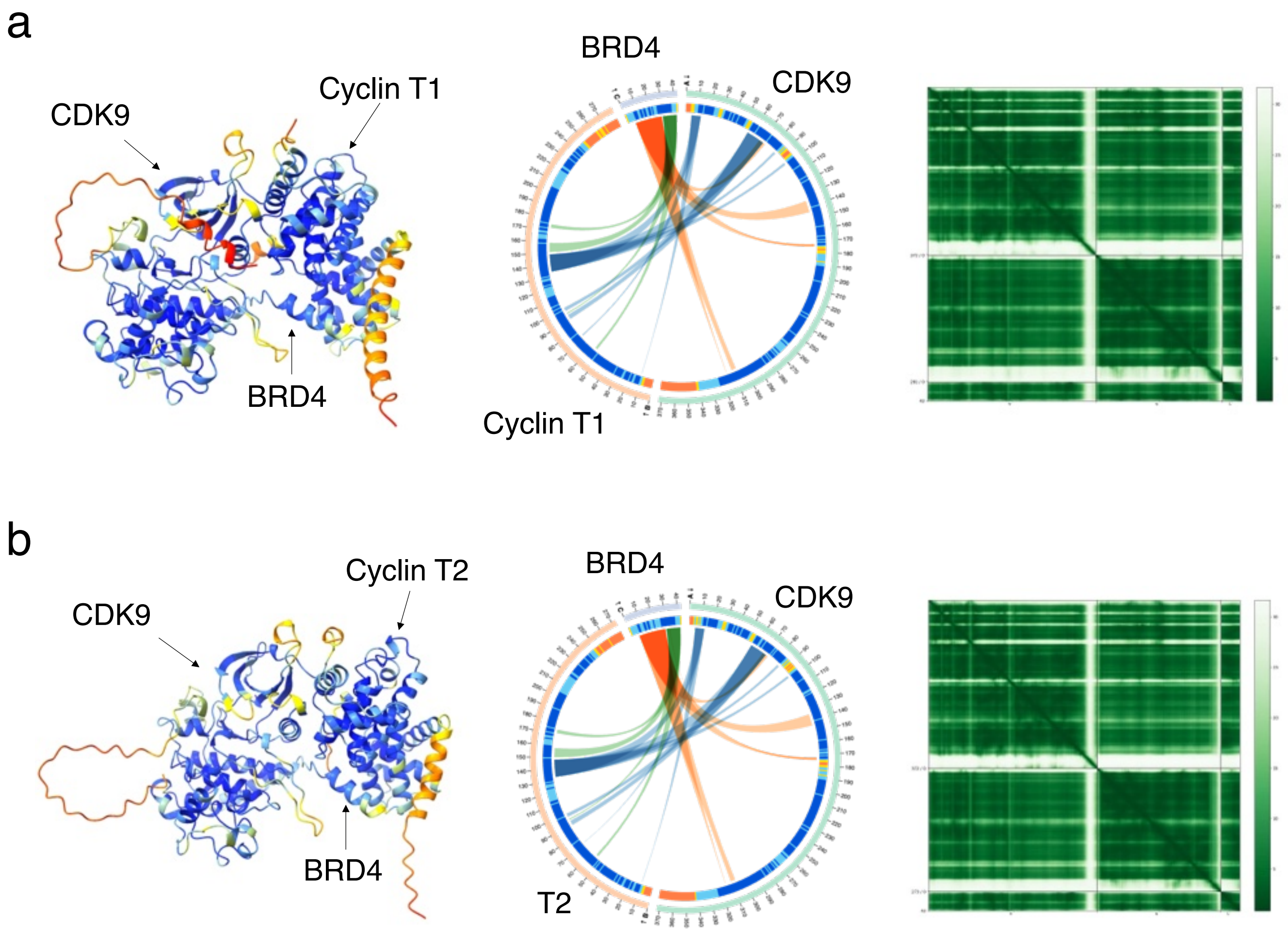

**Supplementary Figure 9. The Alphafold model of the CDK9-cyclin T-BRD4 complex. (a, b)** The AlphaFold3 model [5] for P-TEFb complexes containing isoforms of cyclin T (cyclin T1 (a) and cyclin T2 (b)) with BRD4. The model ribbons are coloured by the predicted Local Distance Difference Test (pLDDT) score (left), the corresponding AlphaBridge plot (middle) [6] and Predicted Aligned Error (PAE) (right) matrices. Structural figures prepared using CCP4mg [3]. Related to Figure 3.

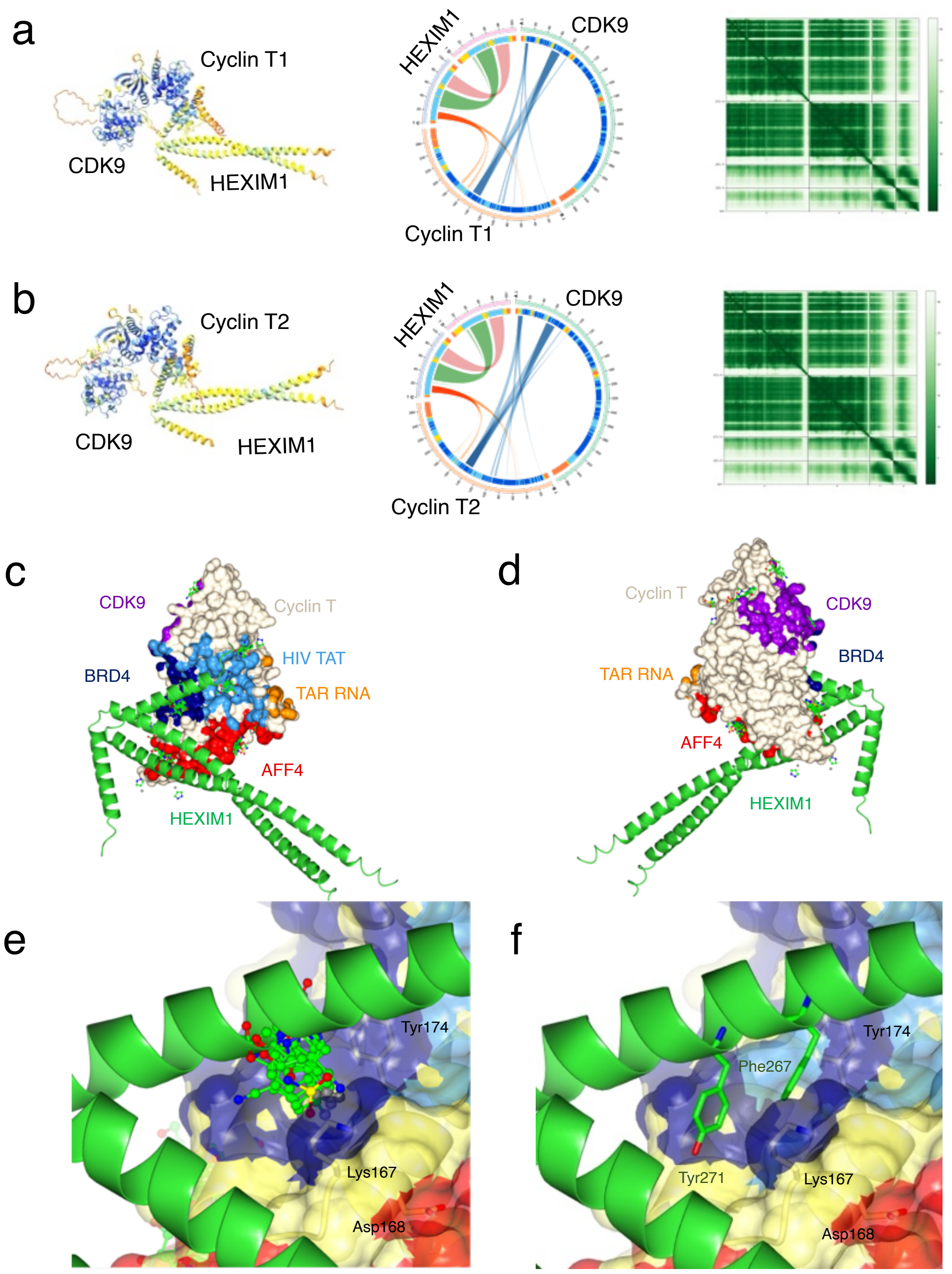

**Supplementary Figure 10. The HEXIM1 binding site on cyclin T. (a, b)** The AlphaFold3 [5] model for P-TEFb complexes containing isoforms of cyclin T1 (a) and cyclin T2 (b) with HEXIM1. The model ribbons are coloured by the predicted Local Distance Difference Test (pLDDT) score (left), the corresponding AlphaBridge plot (middle) [6] and Predicted Aligned Error (PAE) (right) matrices. **(c, d)** Superposition of the HEXIM1 model with the surface of cyclin T1. The cyclin T1 surface occupied by crystallised protein partners (PDB 6CYT) are shown as follows (CDK9, purple; AFF4, red; HIV Tat, blue; TAR, orange). The cyclin T surface that binds to BRD4 as modelled by AlphaFold3 is shown in navy. HEXIM1 (residues Met255-Asp359) is shown in green. (c) and (d) panels are rotated by 180°. **(e, f)** Modelled partner protein HEXIM1 (green ribbon) bound at FragLite Site 3 (the Tat Loop site), overlapping the Tat loop and BRD4 binding site. **(f)** Residues Phe267 and Tyr271 of HEXIM1 are the residues, shown in stick representation, that interact with conserved residues Lys167 ($Lys168_{T1}$), Asp168 ($Asp169_{T1}$), Tyr174 ($Tyr175_{T1}$) on cyclin T2 as identified by mutagenesis experiments as critical to the HEXIM1 interaction [7]. Figure prepared with CCP4mg [3]. Related to Figure 3.

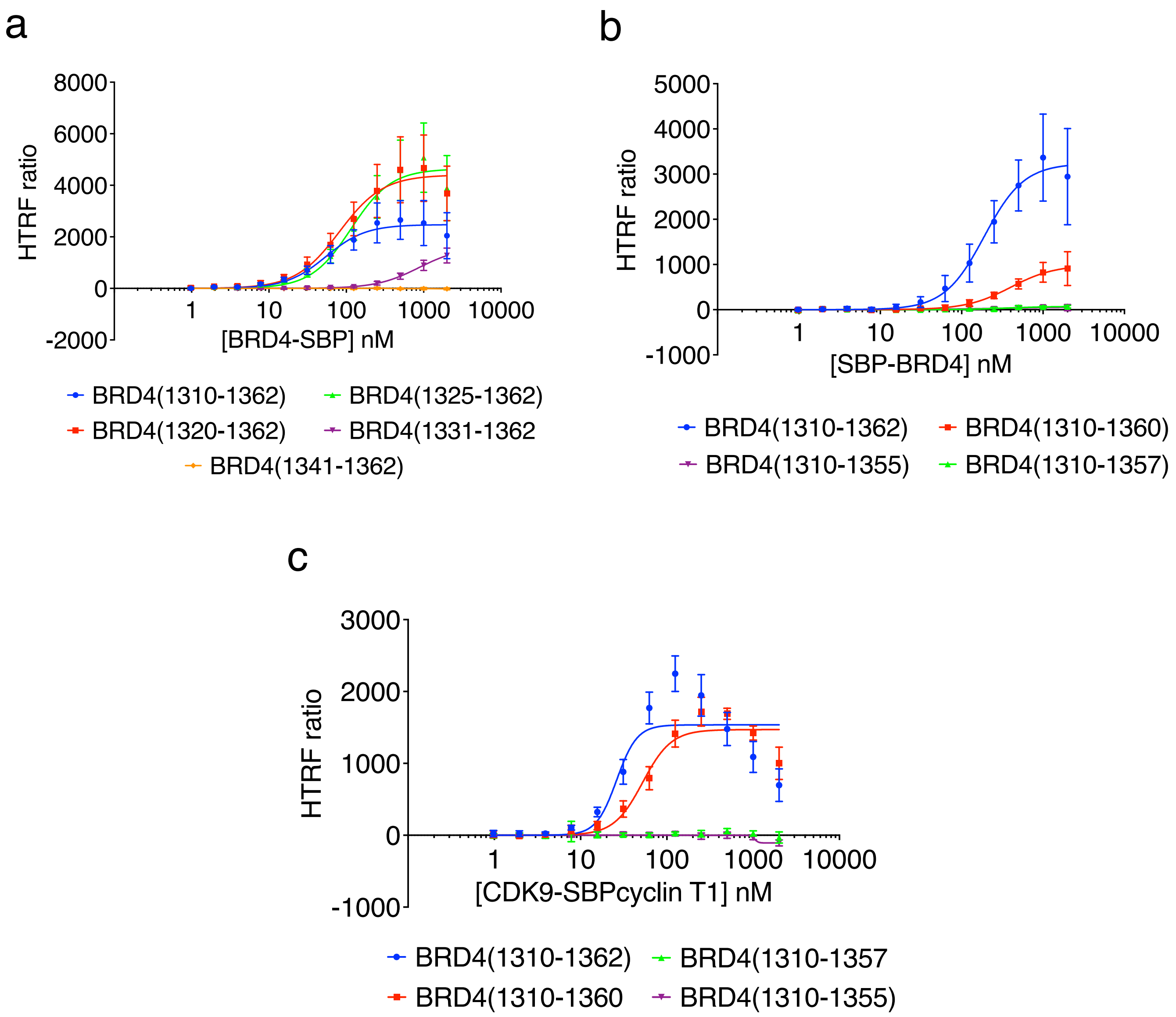

**Supplementary Figure 11. Biophysical characterisation of the interaction between CDK9-cyclin T complexes and BRD4. (a, b)** N-terminally (a) and C-terminally (b) truncated BRD4-SBP and SBP-BRD4 constructs respectively map the termini of the minimal BRD4 P-TEFb Interaction Domain (PID) sequence. The interactions were measured by Homogenous Time-Resolved Fluorescence (HTRF), CDK9-cyclin T1 was labelled with an N-terminal Flag tag. **(c)** HTRF assay to confirm binding of the BRD4 P-TEFb Interaction Domain (PID) construct (residues 1310-1362) to CDK9/SBPcyclin T1. HTRF experiments were carried out in triplicate and repeated on 3 separate days. The error bars indicate SD. Derived Kd values are compiled in Table 1. **(**Related to Figure 4 and Table 1).

**Table S1.** FragLite/PepLite structures, IUPAC names and SMILES string. Related to Figure 1.

| **FragLite** | **Structure** | Name | SMILES |
|---|---|---|---|
| **1** | 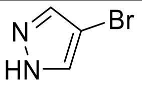 | 4-bromo-1H-pyrazole | BrC1=CNN=C1 |
| **2** | 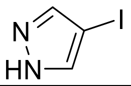 | 4-iodo-1H-pyrazole | IC1=CNN=C1 |
| **3** | 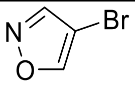 | 4-bromoisoxazole | BrC1=CON=C1 |
| **4** | 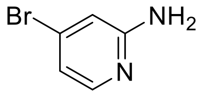 | 4-bromopyridin-2-amine | NC1=NC=CC(Br)=C1 |
| **5** | 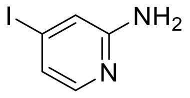 | 4-iodopyridin-2-amine | NC1=NC=CC(I)=C1 |
| **6** | 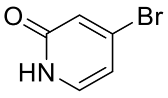 | 4-bromopyridin-2(1H)-one | O=C1C=C(Br)C=CN1 |
| **7** | 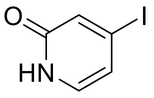 | 4-iodopyridin-2(1H)-one | O=C1C=C(I)C=CN1 |
| **8** | 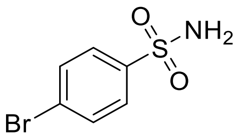 | 4-bromobenzenesulfonamide | BrC1=CC=C(S(N)(=O)=O)C=C1 |
| **9** | 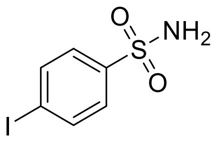 | 4-iodobenzenesulfonamide | IC1=CC=C(S(N)(=O)=O)C=C1 |
| **10** | 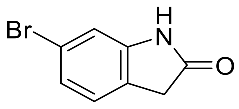 | 6-bromoindolin-2-one | O=C1CC2=C(C=C(Br)C=C2)N1 |
| **11** | 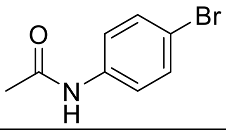 | N-(4-bromophenyl)acetamide | BrC1=CC=C(NC(C)=O)C=C1 |
| **12** | 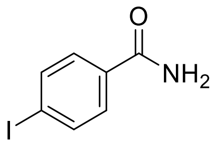 | 4-iodobenzamide | IC1=CC=C(C(N)=O)C=C1 |
| **13** | 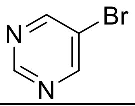 | 5-bromopyrimidine | BrC1=CN=CN=C1 |
| **14** | 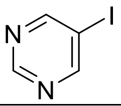 | 5-iodopyrimidine | IC1=CN=CN=C1 |
| **15** | 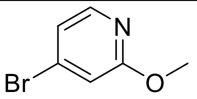 | 4-bromo-2-methoxypyridine | BrC1=CC(OC)=NC=C1 |
| **16** | 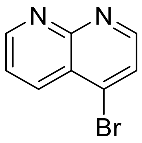 | 4-bromo-1,8-naphthyridine | BrC1=CC=NC2=NC=CC=C21 |
| **17** | 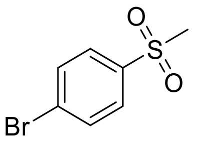 | 1-bromo-4-(methylsulfonyl)benzene | BrC1=CC=C(S(C)(=O)=O)C=C1 |
| **18** | 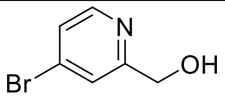 | (4-bromopyridin-2-yl)methanol | BrC1=CC(CO)=NC=C1 |
| **19** | 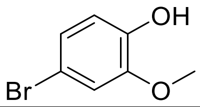 | 4-bromo-2-methoxyphenol | OC1=C(OC)C=C(Br)C=C1 |
| **20** | 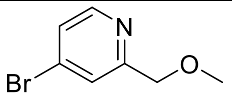 | 4-bromo-2-(methoxymethyl)pyridine | BrC1=CC(COC)=NC=C1 |
| **21** | 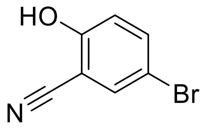 | 5-bromo-2-hydroxybenzonitrile | OC1=C(C#N)C=C(Br)C=C1 |
| **22** | 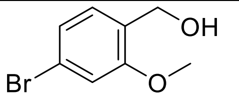 | (4-bromo-2-methoxyphenyl)methanol | BrC1=CC(OC)=C(CO)C=C1 |
| **23** | 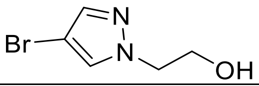 | 2-(4-bromo-1H-pyrazol-1-yl)ethan-1-ol | BrC1=CN(CCO)N=C1 |
| **24** | 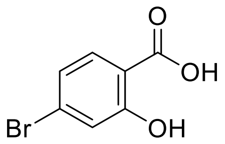 | 4-bromo-2-hydroxybenzoic acid | BrC1=CC(O)=C(C(O)=O)C=C1 |

| | | | |
|---|---|---|---|
| **25** | 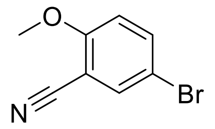 | 5-bromo-2-methoxybenzonitrile | BrC1=CC(C#N)=C(OC)C=C1 |
| **26** | 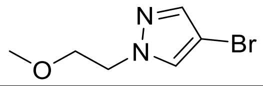 | 4-bromo-1-(2-methoxyethyl)-1H-pyrazole | BrC1=CN(CCOC)N=C1 |
| **27** | 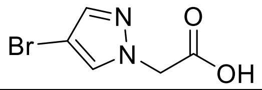 | 2-(4-bromo-1H-pyrazol-1-yl)acetic acid | BrC1=CN(CC(O)=O)N=C1 |
| **28** | 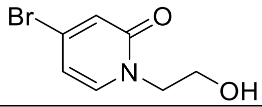 | 4-bromo-1-(2-hydroxyethyl)pyridin-2(1H)-one | O=C1N(CCO)C=CC(Br)=C1 |
| **29** | 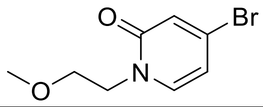 | 4-bromo-1-(2-methoxyethyl)pyridin-2(1H)-one | O=C1N(CCOC)C=CC(Br)=C1 |
| **30** | 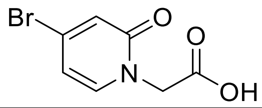 | 2-(4-bromo-2-oxopyridin-1(2H)-yl)acetic acid | O=C1N(CC(O)=O)C=CC(Br)=C1 |
| **31** | 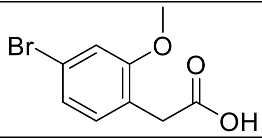 | 2-(4-bromo-2-methoxyphenyl)acetic acid | BrC1=CC(OC)=C(CC(O)=O)C=C1 |
| **PLF** | 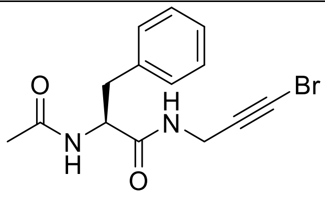 | (S)-2-acetamido-N-(3-bromoprop-2-yn-1-yl)-3-phenylpropanamide | CC(N[C@H](C(NCC#CBr)=O)CC1=CC=CC=C1)=O |
| **PLW** | 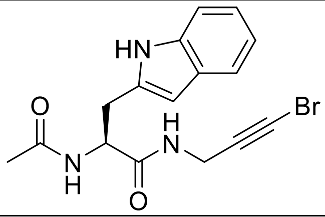 | (S)-2-acetamido-N-(3-bromoprop-2-yn-1-yl)-3-(1H-indol-2-yl)propanamide | CC(N[C@H](C(NCC#CBr)=O)CC1=CC2=C(N1)C=CC=C2)=O |
| **PLY** | 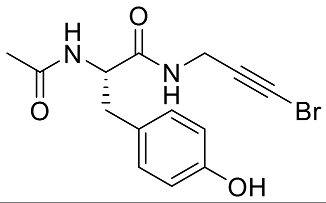 | (S)-2-acetamido-N-(3-bromoprop-2-yn-1-yl)-3-(4-hydroxyphenyl)propanamide | CC(N[C@H](C(NCC#CBr)=O)CC1=CC=C(O)C=C1)=O |
| **PLG** | 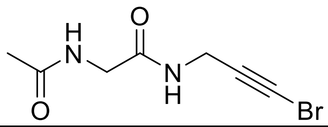 | 2-acetamido-N-(3-bromoprop-2-yn-1-yl)acetamide | CC(NCC(NCC#CBr)=O)=O |
| **PLA** | 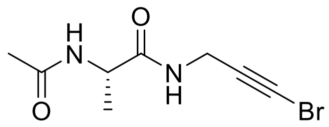 | (S)-2-acetamido-N-(3-bromoprop-2-yn-1-yl)propanamide | CC(N[C@H](C(NCC#CBr)=O)C)=O |
| **PLV** | 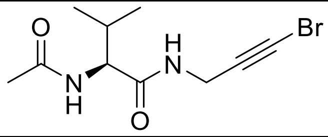 | (S)-2-acetamido-N-(3-bromoprop-2-yn-1-yl)-3-methylbutanamide | CC(N[C@H](C(NCC#CBr)=O)C(C)C)=O |
| **PLL** | 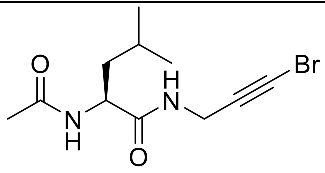 | (S)-2-acetamido-N-(3-bromoprop-2-yn-1-yl)-4-methylpentanamide | CC(N[C@H](C(NCC#CBr)=O)CC(C)C)=O |
| **PLI** | 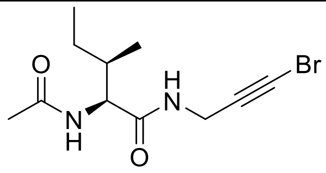 | (2S,3R)-2-acetamido-N-(3-bromoprop-2-yn-1-yl)-3-methylpentanamide | CC(N[C@H](C(NCC#CBr)=O)[C@@H](CC)C)=O |
| **PLM** | 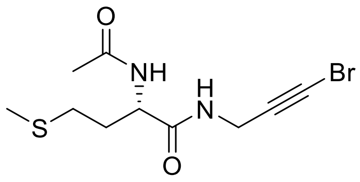 | (S)-2-acetamido-N-(3-bromoprop-2-yn-1-yl)-4-(methylthio)butanamide | CC(N[C@H](C(NCC#CBr)=O)CCSC)=O |
| **PLP** | 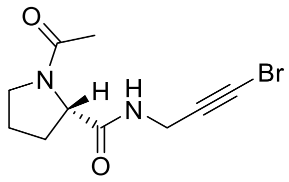 | (S)-1-acetyl-N-(3-bromoprop-2-yn-1-yl)pyrrolidine-2-carboxamide | O=C(NCC#CBr)[C@@]1(CCCN1C(C)=O)[H] |
| **PLS** | 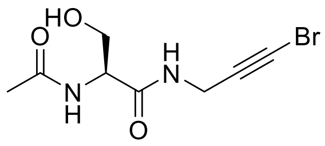 | (S)-2-acetamido-N-(3-bromoprop-2-yn-1-yl)-3-hydroxypropanamide | CC(N[C@H](C(NCC#CBr)=O)CO)=O |
| **PLT** | 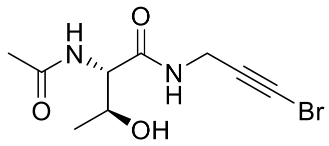 | (2S,3S)-2-acetamido-N-(3-bromoprop-2-yn-1-yl)-3-hydroxybutanamide | CC(N[C@H](C(NCC#CBr)=O)[C@H](C)O)=O |

| | | | |
|---|---|---|---|
| **PLN** | 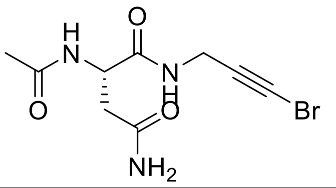 | (S)-2-acetamido-N1-(3-bromoprop-2-yn-1-yl)succinamide | CC(N[C@H](C(NCC#CBr)=O)CC(N)=O)=O |
| **PLQ** | 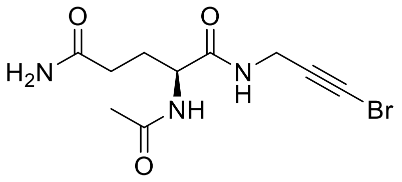 | (S)-2-acetamido-N1-(3-bromoprop-2-yn-1-yl)pentanediamide | CC(N[C@H](C(NCC#CBr)=O)CCC(N)=O)=O |
| **PLD** | 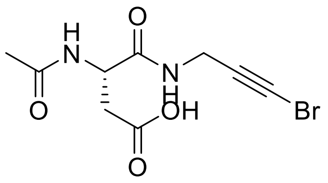 | (S)-3-acetamido-4-((3-bromoprop-2-yn-1-yl)amino)-4-oxobutanoic acid | CC(N[C@H](C(NCC#CBr)=O)CC(O)=O)=O |
| **PLE** | 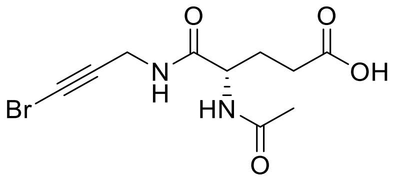 | (S)-4-acetamido-5-((3-bromoprop-2-yn-1-yl)amino)-5-oxopentanoic acid | CC(N[C@H](C(NCC#CBr)=O)CCC(O)=O)=O |
| **PLK** | 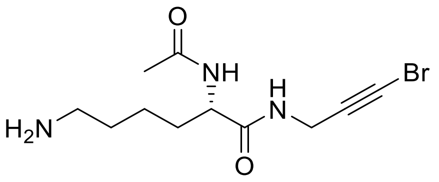 | (S)-2-acetamido-6-amino-N-(3-bromoprop-2-yn-1-yl)hexanamide | CC(N[C@H](C(NCC#CBr)=O)CCCCN)=O |
| **PLR** | 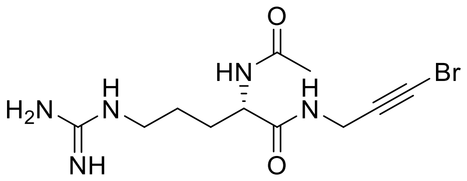 | (S)-2-acetamido-N-(3-bromoprop-2-yn-1-yl)-5-guanidinopentanamide | CC(N[C@H](C(NCC#CBr)=O)CCCNC(N)=N)=O |
| **PLH** | 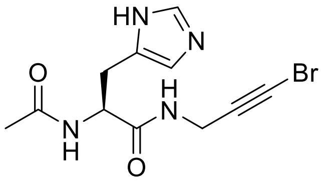 | (S)-2-acetamido-N-(3-bromoprop-2-yn-1-yl)-3-(1H-imidazol-5-yl)propanamide | CC(N[C@H](C(NCC#CBr)=O)CC1=CN=CN1)=O |
| **PLKAc** | 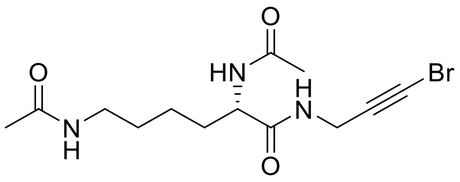 | (S)-N,N'-(6-((3-bromoprop-2-yn-1-yl)amino)-6-oxohexane-1,5-diyl)diacetamide | CC(N[C@H](C(NCC#CBr)=O)CCCCNC(C)=O)=O |

**Table S2. Data statistics and refinement details**

| **Complex** | **CyclinT2-FL1** | **CyclinT2-FL2** | **CyclinT2-FL4** | **CyclinT2-FL5** | **CyclinT2-FL6** | **CyclinT2-FL8** | **CyclinT2-FL9** | **CyclinT2-FL12** |
|---|---|---|---|---|---|---|---|---|
| Beamline | i04-1 | i04-1 | i04-1 | i04-1 | i04-1 | i04-1 | i04-1 | i04-1 |
| PDB code | 9TQV | 9TQW | 9TQX | 9TQZ | 9TQY | 9TR1 | 9TR3 | 9TR4 |
| **Data Collection** | | | | | | | | |
| Wavelength (Å) | 0.916 | 0.916 | 0.916 | 0.916 | 0.916 | 0.916 | 0.916 | 0.916 |
| Space Group | I121 | I121 | I121 | I121 | I121 | I121 | I121 | I121 |
| Unit Cell dimensions (a,b,c)(Å) | 42.0, 59.5, 131 | 136, 59.4, 41.8 | 137, 59.7,42.3 | 136, 59.4, 42.4 | 136, 59.3, 42.4 | 136, 59.7, 42.5 | 42.6, 58.7, 130 | 135.7, 59.2, 42.2 |
| Unit Cell dimensions (α,β,γ)(°) | 90.0, 92.8 90.0 | 90.0, 105, 90.0 | 90.0, 106, 90.0 | 90.0 106, 90.0 | 90.0, 106, 90.0 | 90.0, 106, 90.0 | 90.0, 92.8, 90.0 | 90.0, 106, 90.0 |
| Resolution (Å) | 40.6-1.62 (1.65-1.62) | 65.7-1.80 (1.84-1.80) | 65.6-1.87 (1.92-1.87) | 54.1-1.64 (1.64-1.67) | 39.8-1.62 (1.65-1.62) | 39.8-1.81 (1.85-1.81) | 41.1-2.30 (2.38-2.30) | 53.9-2.10 (2.16-2.10) |
| Rmerge | 0.026 (0.634) | 0.033 (0.535) | 0.05 (0.721) | 0.017 (0.521) | 0.022 (0.608) | 0.031 (0.537) | 0.145 (0.409) | 0.039 (0.460) |
| I/σ(I) | 13.5 (1.1) | 11.7 (1.3) | 8.5 (1.0) | 14.9 (1.2) | 13.9 (1.1) | 12.6 (1.4) | 9.1 (3.9) | 10.6 (1.7) |
| Completeness | 99.8 (98.4) | 100 (99.8) | 99.7 (98.7) | 100 (100) | 99.9 (99.8) | 99.2 (99.0) | 99.6 (98.8) | 98.8 (99.4) |
| CC 1/2 | 0.999 (0.619) | 0.999 (0.645) | 0.997 (0.480) | 1.00 (0.652) | 0.999 (0.633) | 0.999 (0.644) | 0.980 (0.869) | 0.999 (0.629) |
| Multiplicity | 2.0 (2.0) | 2.0 (2.0) | 2.0 (2.0) | 2.0 (2.0) | 2.0 (2.0) | 2.0 (2.0) | 3.8 (3.8) | 1.9 (2.0) |
| Anomalous Completeness | 99.5 (98.3) | 99.7 (99.1) | 99.5 (98.5) | 99.8 (99.9) | 99.7 (99.7) | 98.9 (98.6) | 99.4 (98.7) | 98.4 (99.5) |
| Anomlalous Multiplicity | 1.0 (1.0) | 1.0 (1.0) | 1.0 (1.0) | 1.0 (1.0) | 1.0 (1.0) | 1.0 (1.0) | 1.9 (1.9) | 1.0 (1.0) |
| **Refinement** | | | | | | | | |
| No. reflections all/free | 41176/2135 | 29945/1463 | 26924/1382 | 39975/2000 | 41245/2089 | 29669/1401 | 14514 / 679 | 18708/1007 |
| R-factor/R-free | 0.226/0.229 | 0.232/0.226 | 0.229/0.237 | 0.221/0.221 | 0.222/0.222 | 0.229/0.238 | 0.251/0.248 | 0.219/0.231 |
| Bonds (Å) | 0.0102 | 0.01 | 0.009 | 0.0102 | 0.0107 | 0.0102 | 0.009 | 0.0077 |
| Angles (°) | 1.47 | 1.48 | 1.41 | 1.58 | 1.6 | 1.53 | 1.41 | |
| Number of Ligands | 7 | 10 | 7 | 2 | 3 | 9 | 2 | 1 |

| **Complex** | **CyclinT2-FL18** | **CyclinT2-FL19** | **CyclinT2-FL21** | **CyclinT2-FL22** | **CyclinT2-FL23** | **CyclinT2-FL29** | **CyclinT2-FL31** | **CDK9-Cyclin T2** |
|---|---|---|---|---|---|---|---|---|
| Beamline | i04-1 | i04-1 | i04-1 | i04-1 | i04-1 | i04-1 | i04-1 | i03 |
| PDB code | 9TR0 | 9TQU | 9TQT | 9TR5 | 9TQS | 9TQR | 9TQP | 9TTD |
| **Data Collection** | | | | | | | | |
| Wavelength (Å) | 0.916 | 0.916 | 0.916 | 0.916 | 0.916 | 0.916 | 0.916 | 0.98 |
| Space Group | I121 | I121 | I121 | I121 | I121 | I121 | I121 | P 21 21 21 |
| Unit Cell dimensions (a,b,c)(Å) | 136, 59.1, 42.5 | 136, 59.7, 42.5 | 135, 58.7, 42.6 | 136, 59.5, 42.7 | 136, 59.6, 42.1 | 135, 58.9, 42.2 | 134, 58.8, 42.0 | 100, 155, 186 |
| Unit Cell dimensions (α,β,γ)(°) | 90.0, 105, 90.0 | 90.0, 106, 90.0 | 90.0, 106, 90.0 | 90.0, 106, 90.0 | 90.0, 106, 90.0 | 90.0, 105, 90.0 | 90.0, 105, 90.0 | 90.0, 90.0, 90.0 |
| Resolution (Å) | 50.0-2.29 (2.37-2.29) | 54.3-1.91 (1.95-1.91) | 41.0-2.00 (2.05-2.00) | 65.5-2.02 (2.07-2.02) | 54.3-1.73 (1.76-1.73) | 53.7-1.98 (2.03-1.98) | 64.9-2.03 (2.08-2.03) | 79.6-3.20 (3.30-3.20) |
| Rmerge | 0.013-0.070 | 0.030 (0.432) | 0.041 (0.471) | 0.047 (0.477) | 0.022 (0.546) | 0.033 (0.550) | 0.035 (0.550) | 0.145 (1.14) |
| I/σ(I) | 26.1 (8.8) | 12.2 (1.7) | 11.1 (1.5) | 9.4 (1.5) | 12.0 (1.1) | 11.6 (1.4) | 14.6 (1.5) | 8.4 (1.5) |
| Completeness | 100 (100) | 100 (99.9) | 100 (99.9) | 99.0 (99.8) | 99.9 (99.6) | 99.6 (100) | 99.0 (99.0) | 99.9 (100) |
| CC 1/2 | 1.00 (0.992) | 0.999 (0.685) | 0.998 (0.609) | 0.998 (0.655) | 0.999 (0.715) | 0.998 (0.664) | 0.999 (0.593) | 0.997 (0.668) |
| Redundancy | 1.9 (2.0) | 2.0 (2.0) | 1.9 (2.0) | 1.9 (2.0) | 2.0 (2.0) | 1.9 (2.0) | 1.9 (2.0) | 7.5 (7.3) |
| Anomalous Completeness | 99.9 (99.9) | 99.8 (99.3) | 99.7 (99.6) | 98.4 (98.4) | 99.8 (99.2) | 98.1 (99.4) | 98.1 (98.3) | 99.9 (100) |
| Anomalous Multiplicity | 1.0 (1.0) | 1.0 (1.0) | 1.0 (1.0) | 1.0 (1.0) | 1.0 (1.0) | 1.0 (1.0) | 1.0 (1.0) | 3.9 (3.7) |
| **Refinement** | | | | | | | | |
| No. reflections all/free | 14747 / 754 | 25524 / 1298 | 21856 / 1137 | 21422 / 1071 | 34018 / 1589 | 22288 / 1155 | 20389 / 1063 | 48273 / 2485 |
| R-factor/R-free | 0.223/0.216 | 0.215 / 0.222 | 0.216 / 0.225 | 0.221/0.223 | 0.222 / 0.230 | 0.225/0.229 | 0.219 / 0.221 | 0.203 / 0.226 |
| Bonds (Å) | 0.0078 | 0.0094 | 0.0094 | 0.0084 | 0.0091 | 0.009 | 0.0087 | 0.0072 |
| Angles (°) | 1.44 | 1.49 | 1.37 | 1.42 | 1.47 | 1.44 | 1.4 | 1.68 |
| Number of Ligands | 10 | 7 | 12 | 6 | 7 | 1 | 7 | N/A |

| **Complex** | **CyclinT2-PLG** | **CyclinT2-PLY** | **CyclinT2-PLS** | **CyclinT2-PLT** |
|---|---|---|---|---|
| Beamline | i04-1 | i04-1 | i04-1 | i04-1 |
| PDB code | 9TQO | 9TQN | 9TQL | 9TQM |
| **Data Collection** | | | | |
| Wavelength (Å) | 0.916 | 0.916 | 0.916 | 0.916 |
| Space Group | I121 | I121 | I121 | I121 |
| Unit Cell dimensions (a,b,c)(Å) | 136, 59.4, 42.8 | 136, 59.2, 42.8 | 136, 59.7, 42.3 | 136, 59.6, 42.3 |
| Unit Cell dimensions (α,β,γ)(°) | 90.0, 106, 90.0 | 90.0, 106, 90.0 | 90.0, 106, 90.0 | 90.0, 105, 90.0 |
| Resolution (Å) | 54.1-1.88 (1.92-1.88) | 53.9-1.87 (1.91-1.87) | 65.4-1.87 (1.91-1.87) | 35.22-1.88 (1.92-1.88) |
| Rmerge | 0.030-0.157 | 0.036-0.106 | 0.037-0.200 | 0.022-0.202 |
| I/σ(I) | 15.7 (5.1) | 24.1 (10.5) | 19.1 (5.2) | 17.1 (3.4) |
| Completeness | 97.7 (99.7) | 98.5 (99.5) | 97.1 (97.5) | 99.7 (100) |
| CC 1/2 | 0.997 (0.939) | 0.994 (0.960) | 0.997 (0.903) | 0.999 (0.920) |
| Redundancy | 1.9 (1.9) | 1.9 (1.9) | 1.8 (1.9) | 1.9 (1.9) |
| Anomalous Completeness | 87.6 (96.5) | 90.1 (93.2) | 84.3 (89.6) | 93.8 (95.9) |
| Anomalous Multiplicity | 0.9 (1.0) | 0.9 (1.0) | 0.9 (1.0) | 0.9 (1.0) |
| **Refinement** | | | | |
| No. reflections all/free | 26236 / 1349 | 26579 / 1365 | 26312 / 1356 | 26592 / 1372 |
| R-factor/R-free | 0.189 / 0.189 | 0.190 / 0.195 | 0.188 / 0.181 | 0.184 / 0.187 |
| Bonds (Å) | 0.0114 | 0.0116 | 0.0099 | 0.0109 |
| Angles (°) | 0.172 | 1.78 | 1.61 | 1.61 |
| Number of Ligands | 3 | 1 | 1 | 1 |

Table S3. Summary of FragLite binding events categorised by detection method and associated anomalous signal RMSD values.

| Complex | Total number of anomalous peaks above 4 | PanDDA | PanDDA + Anomalous | Ligand 1 (Peak rmsd) | Ligand 2 | Ligand 3 | Ligand 4 | Ligand 5 | Ligand 6 | Ligand 7 | Ligand 8 | Ligand 9 | Ligand 10 | Ligand 11 | Ligand 12 |
|---|---|---|---|---|---|---|---|---|---|---|---|---|---|---|---|
| **CyclinT2-FL1** | 7 | 2 | 7 | A/301 (41.33) | A/401 (24.90) | A/803 (10.42) | A/810 (8.60) | A/801 (8.18) | A/814 (7.23) | A/811 (6.93) | | | | | |
| **CyclinT2-FL2** | 10 | 4 | 10 | A/301 (16.64) | A/905/A (12.81) | A/905/B (12.81) | A/601/B (7.71) | A/601/C (7.71) | A/501 (5.83) | A/907 (5.24) | A/701 (5.21) | A/903 (4.92) | A/906 (4.50) | | |
| **CyclinT2-FL4** | 7 | 2 | 7 | A/301 (12.01) | A/401 (6.62) | A/403 (6.48) | A/405 (5.49) | A/402 (5.41) | A/406/A (5.35) | A/406/B (5.35) | | | | | |
| **CyclinT2-FL5** | 2 | 1 | 2 | A/301 (10.37) | A/302 (6.48) | | | | | | | | | | |
| **CyclinT2-FL6** | 3 | 1 | 3 | A/301 (36.31) | A/302 (11.10) | A/303 (6.71) | | | | | | | | | |
| **CyclinT2-FL8** | 9 | 3 | 9 | A/301 (29.29) | D/1/A (15.52) | D/1/B (15.52) | D/2/A (9.58) | D/2/B (9.58) | A/401 (7.09) | D/3 (6.94) | D/4 (6.34) | D/5 (5.09) | | | |
| **CyclinT2-FL9** | 2 | 0 | 2 | B/1 (6.18) | B/2 (4.54) | | | | | | | | | | |
| **CyclinT2-FL12** | 1 | 1 | 1 | A/301 (3.88) | | | | | | | | | | | |
| **CyclinT2-FL18** | 10 | 1 | 10 | D/5/A (12.56) | D/5/B (12.56) | D/7/A (12.56) | D/7/B (12.56) | D/4/B (10.70) | D/6 (9.16) | D/8 (8.88) | D/3/A (7.00) | D/3/B (7.00) | D/4/A (4.79) | | |
| **CyclinT2-FL19** | 7 | 3 | 7 | A/301/C (13.89) | A/402 (13.27) | A/403 (9.98) | A/301/B (8.28) | A/404/B (6.54) | A/404/C (5.08) | A/404/A (5.08) | | | | | |
| **CyclinT2-FL21** | 12 | 5 | 12 | A/301 (19.02) | A/501/C (15.76) | A/601/B (9.08) | D/602 (8.86) | A/601/C (8.67) | D/603 (7.94) | D/604 (6.88) | D/605 (6.40) | A/603 (5.34) | A/501/B (5.34) | D/608 (5.00) | D/609 (4.05) |
| **CyclinT2-FL22** | 6 | 3 | 6 | A/301/C (12.62) | A/501 (8.36) | A/401/C (7.77) | A/601 (6.96) | A/301/B (5.15) | A/401/B (4.15) | | | | | | |
| **CyclinT2-FL23** | 7 | 0 | 7 | A/801/A (10.07) | A/801/C (10.07) | A/701/A (8.43) | A/701/B (6.76) | A/802/A (5.90) | A/802/B (5.90) | A/804 (5.54) | | | | | |
| **CyclinT2-FL29** | 1 | 0 | 1 | D/1 (9.07) | | | | | | | | | | | |
| **CyclinT2-FL31** | 7 | 4 | 7 | A/501 (24.29) | A/401 (20.29) | A/301 (12.20) | A/601/C (6.99) | A/604 (6.91) | A/601/A (6.79) | A/605 (6.15) | | | | | |
| **CyclinT2-PLG** | 3 | 3 | 3 | A/301 (14.34) | D/501 (8.66) | A/501 (7.87) | | | | | | | | | |
| **CyclinT2-PLY** | 1 | 1 | 1 | A/401 (14.54) | | | | | | | | | | | |
| **CyclinT2-PLS** | 1 | 1 | 1 | A/301 (18.62) | | | | | | | | | | | |
| **CyclinT2-PLT** | 1 | 1 | 1 | A/401 | | | | | | | | | | | |

**Table S4. Summary of FragLite binding events at partner protein interaction sites.**

| Complex | Total Number of Events | Site 1: AFF4 Loop | Site 2: AFF4 Helix | Site 3: Tat Loop | Site 4: Tat Tail | Site 5: Tat Helix | Site 6 | Site 7: CDK9 N-terminus | Site 8: CDK9 β-strand | Site 9: CDK9 α-Helix | Site 10 | Site 11 | Site 12 | Site 13 | Site 14 |
|---|---|---|---|---|---|---|---|---|---|---|---|---|---|---|---|
| **CyclinT2-FL1** | 7 | A/401 | A/301 | | A/801 | A/803 | A/814 | | | | | | A/810 | A/811 | |
| **CyclinT2-FL2** | 10 | A/905A, A/905/B | A/301 | A/906, A/907 | A/501 | | A/601/B, A/601/C | A/701 | | | | A/903 | | | |
| **CyclinT2-FL4** | 7 | A/403 | A/301 | A/402 | A/406/A, A/406/B | A/401 | | A/405 | | | | | | | |
| **CyclinT2-FL5** | 2 | A/302 | A/301 | | | | | | | | | | | | |
| **CyclinT2-FL6** | 3 | A/302 | A/301 | | | A/303 | | | | | | | | | |
| **CyclinT2-FL8** | 9 | A/301, D/1, D/6 | D2/A, D/2/B | A/401 | D/3 | | D/4 | | | | D/5 | | | | |
| **CyclinT2-FL9** | 2 | B/1, B/2 | | | | | | | | | | | | | |
| **CyclinT2-FL12** | 1 | | A/301 | | | | | | | | | | | | |
| **CyclinT2-FL18** | 10 | D/5/A, D/5/B, D/7/A, D/7/B, D/8 | D/4/A, D/4/B | D/6 | D/3/A, D/3/B | | | | | | | | | | |
| **CyclinT2-FL19** | 7 | A/404/A, A/404/B, A/404/C | A/301, A/401 | A/402 | | A/403 | | | | | | | | | |
| **CyclinT2-FL21** | 12 | D/603/B, D/603/C | A/301 | A/501/B, A/501/C | D/608 | A/601, D/601 | D/602 | D/604 | D/605 | D/609 | | | | | |
| **CyclinT2-FL22** | 6 | A/401/B, A/401/C | A/301/B, A/301/C, A/601 | A/501 | | | | | | | | | | | |
| **CyclinT2-FL23** | 6 | A/801/A, A/801/C | A/701/A, A/701/B | A/802/A, A/802/B | | | | A/804 | | | | | | | |
| **CyclinT2-FL29** | 1 | D/1 | | | | | | | | | | | | | |
| **CyclinT2-FL31** | 7 | A/601/B, A/601/C | A/301 | A/501 | | | A/401 | | A/602/A, A/602/B | | | | | | |
| **CyclinT2-PLG** | 3 | A/501, D/501 | A/301 | | | | | | | | | | | | |
| **CyclinT2-PLY** | 1 | | | A/401 | | | | | | | | | | | |
| **CyclinT2-PLS** | 1 | | | | | | | | | | | | | | A/301 |
| **CyclinT2-PLT** | 1 | | | | | | | | | | | | | | A/401 |
| | | 30 | 20 | 13 | 8 | 6 | 6 | 4 | 3 | 1 | 1 | 1 | 1 | 1 | 2 |

**KEY RESOURCES TABLE**

| REAGENT or RESOURCE | SOURCE | IDENTIFIER |
|---|---|---|
| **Antibodies** | | |
| Tb-anti-Flag antibody | CisBio | 61FG2TLF |
| **Bacterial and Virus Strains** | | |
| *E. coli* DH5α | Invitrogen | 18265-017 |
| *E. coli* Rosetta (DE3) | Novagen | 70954 |
| *E. coli* Rosetta2 (DE3) pLysS | Novagen | 71403 |
| *E. coli* DH10EmBacY | Geneva Biotech | Supplied as part of MultiBac™ Expression System Kit |
| **Chemicals, Peptides, and Recombinant Proteins** | | |
| FragLite chemical library (Supplementary Table S1) | CancerTools.org Wood et al. [8] | 155084 |
| Glutathione resin | Cytiva | 17075605 |
| 5 mL GSTrap FF column | Cytiva | 17513102 |
| LB Broth | | |
| Protease inhibitor cocktail tablets | Roche | 40694200 |
| GeneJuice® Transfection reagent | Merck Millipore | 70967 |
| Streptavidin-labelled XL665 | CisBio | 610SAXLB |
| MultiBac™ Expression System Kit | Geneva Biotech | N/A |
| Insect-Xpress insect cells medium | Lonza | BELN12-730Q |
| **Critical Commercial Assays** | | |
| Coomassie Protein Assay Reagent | Pierce | 23238 |
| **Deposited Data** | | |
| Cyclin T2-FL1 | 9TQV | This paper |
| Cyclin T2-FL2 | 9TQW | This paper |
| Cyclin T2-FL4 | 9TQX | This paper |
| Cyclin T2-FL5 | 9TQZ | This paper |
| Cyclin T2-FL6 | 9TQY | This paper |
| Cyclin T2-FL8 | 9TR1 | This paper |
| Cyclin T2-FL9 | 9TR3 | This paper |
| Cyclin T2-FL12 | 9TR4 | This paper |
| Cyclin T2-FL18 | 9TR0 | This paper |
| Cyclin T2-FL19 | 9TQU | This paper |

| | | |
|---|---|---|
| Cyclin T2-FL21 | 9TQT | This paper |
| Cyclin T2-FL22 | 9TR5 | This paper |
| Cyclin T2-FL23 | 9TQS | This paper |
| Cyclin T2-FL29 | 9TQR | This paper |
| Cyclin T2FL31 | 9TQP | This paper |
| CDK9-cyclin T2 | 9TTD | This paper |
| Cyclin T2-PLG | 9TQO | This paper |
| Cyclin T2-PLY | 9TQN | This paper |
| Cyclin T2-PLS | 9TQL | This paper |
| Cyclin T2-PLT | 9TQM | This paper |
| **PDB codes for structures used within, but not derived from, this study** | | |
| CDK9-cyclin T1 | 3BLH | [9] |
| CDK9-cyclin T1-AFF4 | 4IMY | [10] |
| CDK9-cyclin T1-HIVTat | 3MI9 | [11] |
| CDK9-cyclin T1-AFF4-HIVTat | 4OGR | [12] |
| CDK9-cyclin T1-AFF4-HIVTat-TAR | 6CYT | [13] |
| Cyclin T2 | 2IVX | [9] |
| **Experimental models: cell lines** | | |
| *Spodoptera frugiperda* Sf9 cells | OET | 600100 |
| **Recombinant DNA** | | |
| BRD4-CTR (1279-1362) | Top Gene Technologies | N/A |
| pET3d-GST-3c-BRD4(1310-1362)-SBP | This paper | N/A |
| pET3d-GST-3c-BRD4(1320-1362)-SBP | This paper | N/A |
| pET3d-GST-3c-BRD4(1325-1362)-SBP | This paper | N/A |
| pET3d-GST-3c-BRD4(1331-1362)-SBP | This paper | N/A |
| pET3d-GST-3c-BRD4(1341-1362)-SBP | This paper | N/A |
| pET3d-GST-3c-SBP-BRD4(1310-1362) | This paper | N/A |
| pET3d-GST-3c-SBP-BRD4(1310-1360) | This paper | N/A |
| pET3d-GST-3c-SBP-BRD4(1310-1357) | This paper | N/A |
| pET3d-GST-3c-SBP-BRD4(1310-1355) | This paper | N/A |
| pET3d-GST-3c-FLAG-BRD4(1310-1362) | This paper | N/A |
| pET3d-GST-3c-FLAG-BRD4(1310-1360) | This paper | N/A |
| pET3d-GST-3c-FLAG-BRD4(1310-1357) | This paper | N/A |
| pET3d-GST-3c-FLAG-BRD4(1310-1355) | This paper | N/A |
| pET3d-GST-3c-BRD4-SBP(1310-1362) Gln1350Ala | This paper | N/A |

| | | |
|---|---|---|
| pET3d-GST-3c-BRD4-SBP(1310-1362) Leu1354Ala | This paper | N/A |
| pET3d-GST-3c-BRD4-SBP(1310-1362) Phe1357Ala | This paper | N/A |
| GST3cBRD4(1310-1362)GFP | | |
| GST3cBRD4(1310-1362)GFP (Phe1349Ala) | | |
| GST3cBRD4(1310-1362)GFP (Gln1350Ala) | | |
| GST3cBRD4(1310-1362)GFP (Leu1354Ala) | | |
| GST3cBRD4(1310-1362)GFP (Phe1357Ala) | | |
| pNIC28-Bsa4 HisTEVcyclin T2 (2-273, Gln130Arg) | Structural Genomics Consortium Oxford | Backbone SGC: https://www.thesgc.org/reagents/toronto/vectors |
| pNIC28-Bsa4 HisTEVcyclin T2 (2-273, Gln130Arg, Tyr174Ala) | This paper | |
| pNIC28-Bsa4 HisTEVcyclin T2 (2-273, Gln130Arg, Phe175Ala) | This paper | |
| pNIC28-Bsa4 HisTEVcyclin T2 (2-273, Gln130Arg, Trp206Ala) | This paper | |
| pET3d GST3cBRD4(1310-1362) | This paper | |
| pET3d His3cSBPcyclinT1 (1-281) | This paper | |
| pACEBAC1 CDK9(1-330)His | This paper | |
| pET3d His3cFlagCyclinT1(1-281) | This paper | |
| pET3d His3cFlagCyclinT1(1-281) Tyr175Ala | This paper | |
| pET3d His3cFlagCyclinT1(1-281) Phe176Ala | This paper | |
| pET3d His3cFlagCyclinT1(1-281) Trp210Ala | This paper | |
| **Software and Algorithms** | | |
| AutoProc | Vonrhein et al [14] | |
| Xia2 and DIALS | Winter et al [15] | |
| CCP4i2 software suite | Potterton et al [16] | |
| Phaser | McCoy et al [17] | |
| COOT | Casañal, A. et al [18] | |
| REFMAC5 | Murschudov et al [19] | |
| CCP4MG | McNicholas et al [3] | |
| AlphaFold 3 | Abramson et al [20] | |

| | | |
|---|---|---|
| AKTA Unicorn 7 Evaluation software | Cytiva | https://www.cytivalifesciences.com/ |
| MicroCal PEAQ Evaluation software | MicoCal | N/A |
| GraphPad Prism | GraphPad Software Inc | https://www.graphpad.com |
| Jalview | Waterhouse et al. [2] | www.jalview.org |
| **Other** | | |
| Insect-XPRESS$^{TM}$ Protein-free Insect Cell Medium | Lonza | 12-730Q |
| MicroCal PEAQ | Malvern Instruments | N/A |
| PHERAstar FS | BMGLabtech | N/A |
| PHERAstar FS excitation filter (337 nm) | BMG Lab tech | N/A |
| PHERAstar FS emission filer (620/665 nm) | BMG Lab tech | N/A |
| PHERAstar FS excitation filter (480 nm) | BMG Labtech | N/A |
| PHERAstar FS emission filter (520 nm) | BMG Labtech | N/A |
| Mosquito robot | SPT Labtech | Bespoke specification |
| Biomek liquid handler | Beckman Coulter | Bespoke specification |
| ECHO acoustic liquid dispenser | Beckman Coulter | Bespoke specification |
| 96 well plate, IMP@CT<br>Clear, for microbatch | Greiner Bio-One | 673101 |
| MRC 2Lens Crystallisation Plate | SWISSCI | UVXPO-2LENS |
| AKTA$^{TM}$ Pure chromatography system | Cytiva | Bespoke specification |

**Supplementary Information References**
